# EWS-FLI1 Expression in Human Embryonic MSCs Leads to Transcriptional Reprograming, Defective DNA Damage Repair and Ewing Sarcoma

**DOI:** 10.1101/2025.01.13.632712

**Authors:** Inmaculada Hernández-Muñoz, Irene Cuervas, Estela Prada, Julian Pulecio, Ramón Gimeno, Evelyn Andrades, Soledad Gómez-González, Pau Berenguer-Molins, Ariadna Acedo-Terrades, Júlia Perera-Bel, Marta Bódalo, Lara Nonell, Elena Pérez, Daniela Grases, Caterina Mata, José Yélamos, Yvonne Richaud, Enrique Vidal, Yasmina Cuartero, François Le Dily, Mariona Suñol, Alejandro Manzanares, Angel Raya, Jaume Mora

## Abstract

Ewing sarcoma (ES) is an aggressive bone and soft tissue neoplasm characterized by *EWSR1/ETS* rearrangements and whose cellular origin remains unclear. EWS-FLI1 expression in human pediatric mesenchymal stem cells (hpMSCs) induces a quantitatively and qualitatively different transcriptional response than its expression in human adult MSCs (haMSCs), but fails to form tumors *in vivo*. ES cells have early developmental lineage signatures distinct from postnatal MSCs. Here, we have generated MSCs from experimental teratomas out of human embryonic stem cells (heSCs). Transduction of these human embryonic mesenchymal stem cells (heMSCs) with EWS-FLI1 results in the acquisition of an ES transcriptome, although the oncogene does not preferentially bind to promoters, but to intronic and intergenic microsatellites with >10 CA dinucleotides and GGAA repeats, respectively. In heMSCs, EWS-FLI1 directly regulates BRCA1 expression, although EWS-FLI1-expressing cells show defects in DNA damage repair. Xenografting of EWS-FLI1-transduced heMSCs resulted in the formation of tumors expressing characteristic ES markers. In summary, EWS-FLI1 enforces an aberrant transcriptome and endows *in vivo* transforming capacity when expressed in an undifferentiated early heMSC. Our approach represents an innovative experimental method for understanding critical aspects of the biology of developmental tumors, from leukemia to sarcomas, in which few (even single) genetic alterations are able to transform a fetal stem cell.

## MAIN

Ewing’s sarcoma family of tumors (ESFT) are rare and highly aggressive malignant bone and soft tissue tumors that mostly affect adolescents and young adults (https://rarediseases.org/rare-diseases/ewing-sarcoma/). ESFTs are characterized by *EWSR1/ETS* translocations, frequently involving *EWSR1* and *FLI1* (hereby *EWS-FLI1*), that appears to be the only detectable oncogenic event in as many as 20% of tumors ^1–3^. Since the initial description of a new entity called “diffuse endothelioma of the bone” by Dr. James Ewing in 1921 ^4^ and the discovery of the fusion oncoprotein in 1992 ^5^, the fundamental question in ESFT research has been to elucidate the identity of the progenitor cell that gives rise to a tumor with a transcriptional profile characteristic of Ewing sarcoma (ES). Current hypothesis is that this rare cancer arises from a stem cell precursor ^6^. Indeed, among different primary human cells, only pluripotent stem cells display permissiveness for exogenous EWS-FLI1 expression ^7,8^. In addition, ES tumors display transcriptional features of mesenchymal stem cells (MSCs)^9^, neural crest stem cells (NCSCs)^10^ and endothelial tissues ^11^. Furthermore, the normal tissues which show Ewing-like transcriptomes belong to developmental transitions such as gastrulation ^12^. These data led to postulate that a primitive neural crest-derived progenitor at the transition to mesenchymal and/or endothelial differentiation could be transformed in ESFTs ^11^.

Human MSCs (hMSCs) are multipotent precursors that can be differentiated *in vitro* into various cell types, typically osteoblasts, chondrocytes, and adipocytes, but also into endothelial lineage ^13^. EWS-FLI1 expression in adult hMSCs (haMSCs) blocks their differentiation, and generates a transcriptome profile reminiscent of ESFT ^8^. Many of these EWS-FLI1-regulated genes are even more induced when the oncogene is expressed in human pediatric MSCs (hpMSCs), while numerous genes that are among the most prominent ESFT markers are induced in hpMSCs but not in haMSCs ^14^. Notably, although hpMSCs provide a far more permissive environment, hpMSCs that express EWS-FLI1 (EF-hpMSCs) are, like EF-haMSCs, unable to form tumors *in vivo*, which has led to the suggestion that a cooperating genetic event or a unique cellular phenotype is additionally required. To address these two issues, Gordon and colleagues developed an approach to model Ewing sarcoma that exploits the *in vitro* potential of human embryonic stem cells (heSCs) to differentiate into cells from the three germ layers when forming embryoid bodies (EBs)^15^. EWS-FLI1 expression in EBs derived from p53-deficient heSCs leads to *in vitro* transformation, yet EB-derived cells lack tumor formation capacities *in vivo* ^15^. It should be noted that although *p53* mutation has been described in up to 50% of ESFT cell lines, *p53* is rarely mutated in primary ES tumors.

The difficulties in identifying a suitable cellular environment that simultaneously recapitulates the oncogene’s ability to block cell differentiation and induce tumor transformation have hindered the understanding of the underlying mechanisms that trigger ESFT and the generation of reliable experimental murine models. Regarding the latter, EWS-FLI1 fails to transform immortalized mouse fibroblasts other than NIH3T3 ^16^, but mouse MSCs transfected with EWS-FLI1 form tumors when injected subcutaneously into nude mice ^7^. More recently, it was shown that *ex vivo* expression of EWS-FLI1 efficiently transforms embryonic osteochondrogenic progenitors, indicating that embryonic precursor cell plasticity may be critical for uncontrolled cell growth ^17^. Despite this tumorigenic ability, the transcriptional profile of the tumors generated by EWS-FLI1 expression in mouse cells only partially resembles the human ESFT transcriptome. Besides, all efforts to model a transgenic mouse that generate ES-like tumors have failed ^18^.

Here we have isolated, cultured and characterized human embryonic mesenchymal stem cells (heMSCs) from heSC-derived experimental teratomas. To investigate whether EWS-FLI1 expression is tolerated, heMSCs were transduced with a Flag-tagged EWS-FLI1 lentivirus. Oncogene expression induced a transcriptional profile indicative of an aberrant differentiation towards an endothelial-neural hybrid phenotype consistent with the ESFT signature. Chromatin immunoprecipitation sequencing (ChIP-seq) revealed that, in addition to intergenic regions, EWS-FLI1 binds mainly to intronic microsatellites (>10 CA dinucleotides) in heMSCs. In addition, we identified *BRCA1*, an essential protein in the DNA repair process, as a direct EWS-FLI1 target. An intrinsic DNA damage repair (DDR) defect characterizes EWS-FLI1-infected heMSCs providing high sensitivity to DNA damaging agents, a characteristic feature of human ES. Inoculation of the transduced cells into the gastrocnemius of mice led to the development of tumour masses in the spine or abdominal cavity as well as lung, liver and kidney metastases. Analysis by spatial transcriptomics of these tumors confirmed a transcriptome compatible with ES and revealed the expression of tumor-specific proteins never described before. In teratoma-derived heMSCs, EWS-FLI1 expression is the only genetic event required to induce ES. This experimental approach provides a valuable platform to explore developmental cancer biology.

## RESULTS

### Teratoma-derived Human Embryonic Mesenchymal Stem Cells (heMSCs) are Earlier Progenitors than hpMSCs

ESFT may arise from early pluripotent precursor cells that tolerate EWS-FLI1 expression, in which the fusion oncoprotein inhibits cell differentiation. Recently, experimental teratomas have been proposed as a useful multilineage model of human development, as they can produce a wide range of cell types from the major developmental lineages transcriptionally similar to the corresponding human fetal cell types ^19^. We used this model as a source of early progenitors in which to assess the effects of EWS-FLI1. As expected, inoculation of heSCs into mouse testes resulted in the formation of teratomas (Figure S1A), which contained elements of the three germ layers (ectoderm, mesoderm and endoderm) (Figure S1B). To isolate progenitor cells, teratomas were disaggregated and adherent cells maintained in culture. Isolated cells exhibited a fibroblastic morphology and human nuclear staining (Figure 1A). Serial culture passages showed that proliferation of heMSCs decreased progressively, and cells eventually stopped proliferating, and became senescent (Figure 1B). heMSCs expressed CD73, CD105, CD90, and HLA-I; lacked CD45, CD34, and CD31; and were able to differentiate to osteogenic, chondrogenic, and adipogenic precursors (Figures 1C, 1D). Therefore, the expression profiling of surface markers and differentiation potential displayed by heMSCs fulfilled the definition of MSCs ^20^.

**Figure 1.**
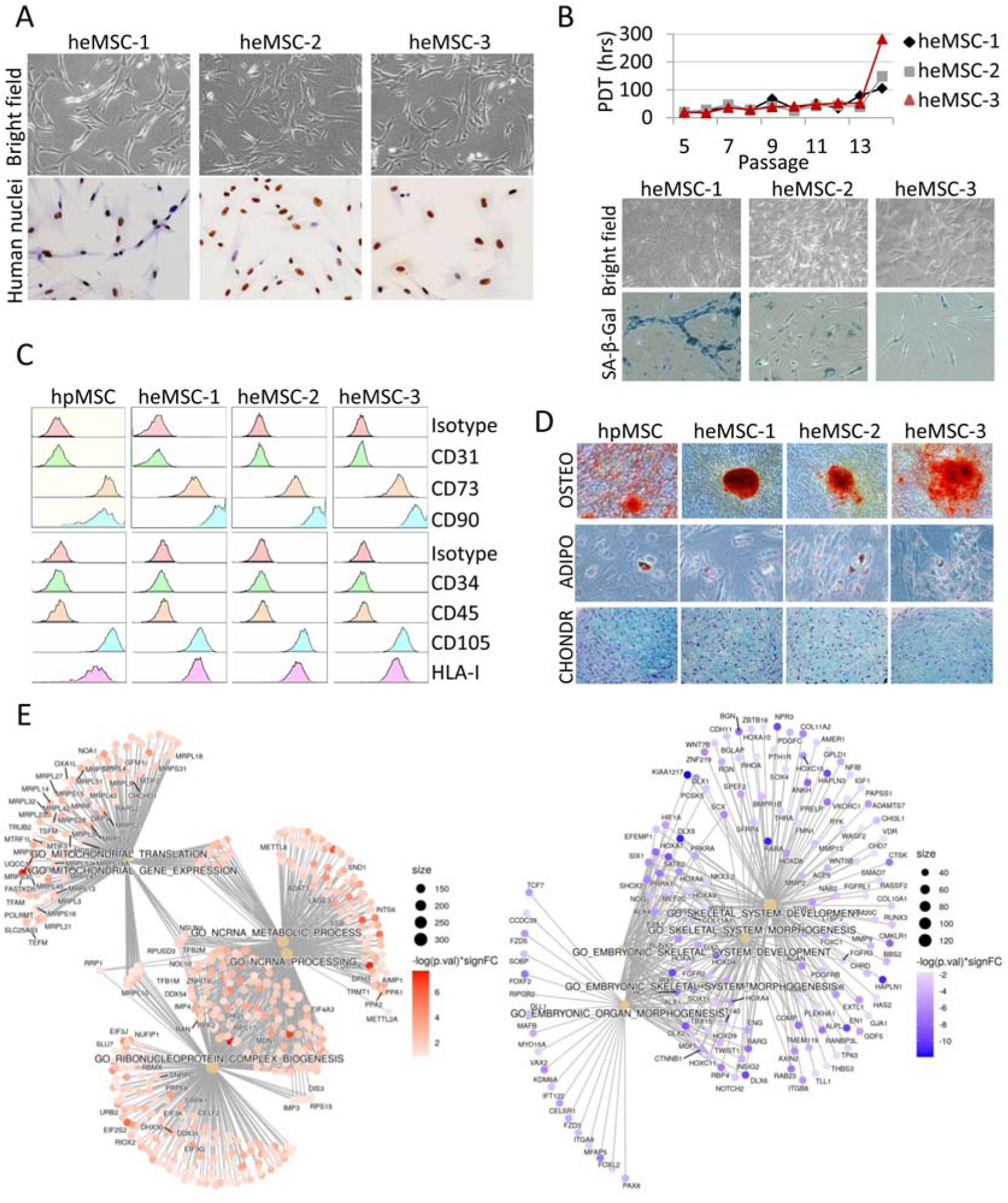
Isolation, Culture and Characterization of heMSCs derived from Experimental Teratomas. (A) Phase contrast microscopy and human nuclei immunohistochemistry of heMSC cultures. (B) Upper panel, proliferative exhaustion of heMSCs during passaging. Data from one representative experiment (from two independent experiments) are presented by plotting population doubling time (PDT) against passages. Lower panel, bright field pictures of heMSCs cultured for more than 20 passages and senescence-associated β-Galactosidase (SA-β-Gal) staining of these cultures. (C) Characterization of surface markers in heMSC by flow cytometry. (D) heMSC differentiation to osteogenic (alizarin red staining), adipogenic (oil red staining) and chondrogenic (alcian blue staining) lineages upon culture with differentiation media. In both C and D, pediatric hMSCs (hpMSCs) are shown as positive control. (E) Gene-concept networks of top 5 significantly enriched terms in control heMSC-1 when compared to hpMSCs, based on the RNA-seq data analysis (adjusted p-value <0.05).

The comparison between heMSCs with hpMSCs isolated from bone marrow aspirates showed that heMSC transcriptomes were enriched in genes specifically involved in RNA processing, while hpMSCs differentially expressed transcripts mainly related to osteochondral specification (Table S1; Figures 1E, S1C). These results clearly reflect the earlier progenitor status of heMSCs compared to hpMSCs.

### EWS-FLI1 Expression in heMSCs Induces a Transcriptional Profile Indicative of ESFT

We investigated the possibility that EWS-FLI1 might rewire the transcriptome of heMSCs more efficiently compared to hpMSCs. Expression levels of the Flag-tagged EWS-FLI1 were significantly lower than the ES cell line A673 and did not affect constitutive expression of the ES marker CD99 (Figure 2A). However, they were sufficient to trigger cell cycle arrest by inducing the expression of the tumor suppressors RB and p53 -and its target gene p21-(Figure 2B), indicating that p53-p21-RB1 signalling is functionally intact in heMSCs. Besides the antiproliferative signal, EWS-FLI1 in heMSCs induces or represses the expression of well-described oncogene targets such as IGF1, LOX and ENC1, as shown in Figure 2C.

**Figure 2.**
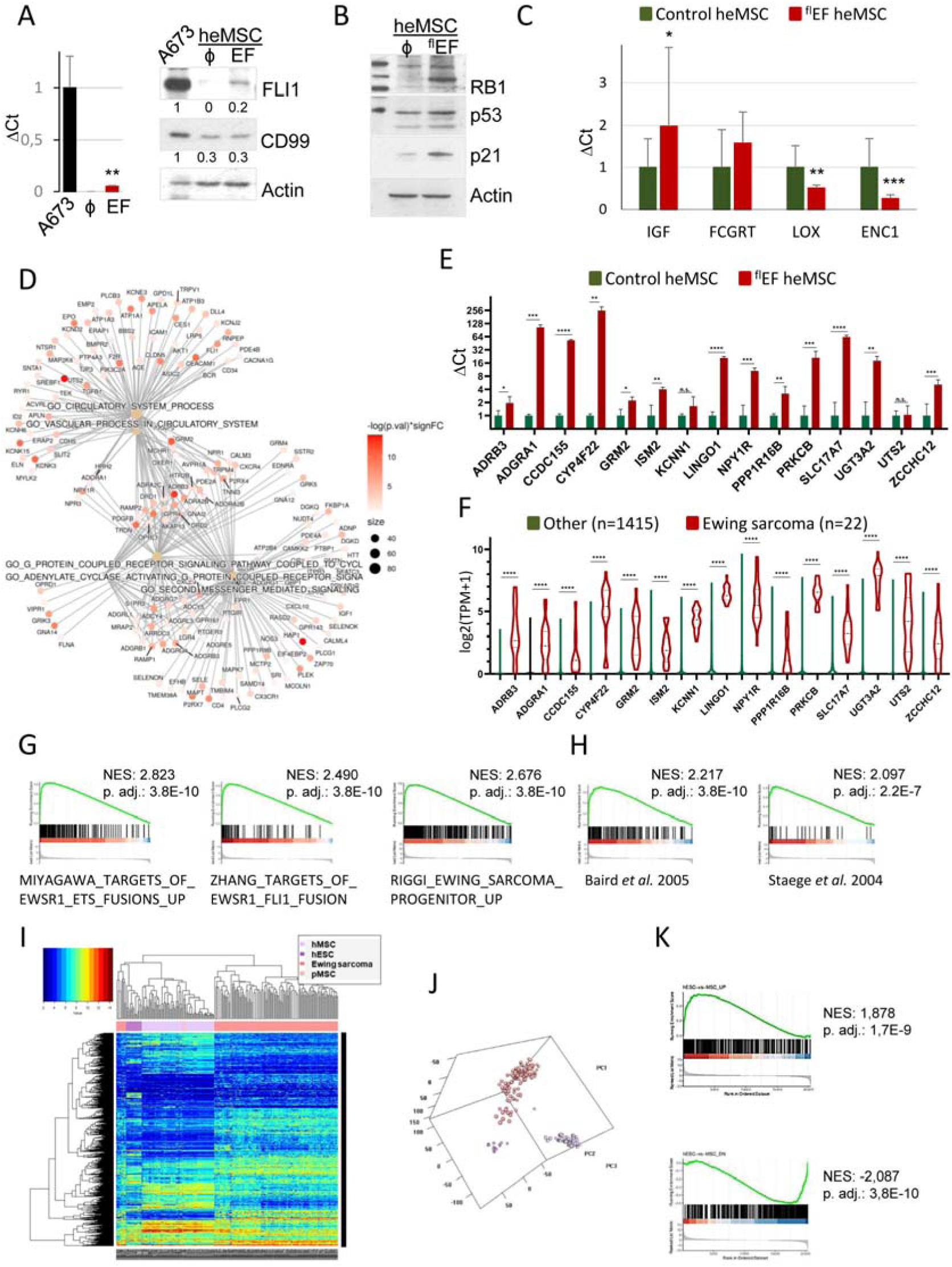
Transcriptional Changes in heMSCs expressing EWS-FLI1 (EF-heMSCs). (A) Expression of EWS-FLI1 and CD99 in heMSCs at 48 hours after infection and A673 cells, as determined by RT-qPCR and Western blot. Actin, loading control. At the bottom, densitometric quantification of Western blot signals, normalized with respect to Actin intensity. (B) Expression of proteins of the p53-p21-RB1 axis in heMSCs at 48 hours after infection with EWS-FLI1, as determined by Western blot. Actin, loading control. (C) Expression of some of the induced and repressed oncogene targets in heMSC-1 cells at 48 hours after infection with EWS-FLI1, as determined by RT-qPCR, and referred to GAPDH mRNA. Data obtained from two independent experiments performed in triplicate. Statistics performed by the Student’s t-test. (D) Gene-concept networks of top 5 significantly enriched terms in EF-heMSCs *vs.* control heMSCs, based on the RNA-seq data analysis (adjusted p-value <0.05). (E) Expression of 15 of the 30 top genes most potently induced by the oncogene, determined by RT-qPCR and referred to GAPDH, in heMSC-1 cells at 48 hours after infection with EWS-FLI1. Data obtained from three independent experiments performed in triplicate. T-Student test was performed. (F) Expression values obtained from DepMap of the genes shown in (E) in Ewing sarcoma and other cancer cell lines (log2(TPM+1)). Mann-Whitney U test was performed. (G) Gene set enrichment analysis (GSEA) of EF-heMSCs transcriptomes in various EWS-FLI1 target signatures: genes commonly up-regulated in UET-13 cells (mesenchymal progenitors) by expression of EWS-FLI1 ^22^; tetracycline-inducible EWS-FLI1 expression system in the rhabdomyosarcoma cell line RD ^10^; and expression of EWS-FLI1 in hMSCs ^8^. (H) GSEA of EF-heMSCs transcriptomes in various Ewing sarcoma signatures ^11,23^. (I) Heatmap and (J) PCA representation of the unsupervised clustering analysis of gene expression signatures in heSCs, hMSCs and Ewing sarcoma samples. Data were obtained from GSE6460, GSE7637, GSE7896, GSE8884, GSE9451, GSE9440, GSE9510, GSE9520, GSE9593, GSE10315, GSE13604, GSE13828, GSE17679, GSE34620, GSE37371 and GSE31215. (K) GSEA of EF-heMSCs transcriptomes in the 400 most expressed and under-represented genes identified by the unsupervised clustering analysis of Ewing sarcomas. P values, *<0.05; **<0.01; ***<0.001; ****<0.0001.

RNA-sequencing and independent analysis of the changes induced by EWS-FLI1 expression in three different heMSC lines (EF-heMSCs) identified 3836 differential expressed genes (DEGs; 2204 and 1632 upregulated and downregulated, respectively; p <0.05) (Table S2; Figure S2A). Normalized enrichment score (NES) analysis of the DEGs revealed that EWS-FLI1 induced transcriptional changes in genes mainly involved in two specific and interconnected functions: signal transduction cascades of receptors coupled to G-proteins (GPCRs) and biological processes involving the circulatory system (Figure 2D; Figure S2B). In contrast, the repressed functions associated with EWS-FLI1 expression in heMSCs were restricted to autophagocytosis and tRNA methylation (Figure S2C). In addition, and in agreement with the neuroectodermal and endothelial features of ESFTs, EF-heMSCs displayed an enhanced expression of proteins involved in signal transduction cascades in neural cells (i.e. NGFR, ALK, NTRK1, RET, GRM2 and NPY1R) and endothelial cells (i.e. UTS2, ACE, CDH5, DLL4, ECE1 or the Angiopoietin-1 receptor TEK). Importantly, most transcriptionally induced genes are non-described EWS-FLI1 targets. A selection of 15 of the 30 genes most potently induced by the oncogene were validated by RT-qPCR (Figure 2E). Expression data extracted from DepMap (https://depmap.org/portal/) show that many of these genes exhibit an expression pattern specific to ES cell lines (Figure 2F). Accordingly, among the most induced genes was *PRKCB*, described by Surdez *et al.* as critical for tumor cell survival and development in ES cells and expressed in primary ES tumors but not in other tumor types ^21^ (Figure S3A).

To estimate further the significance of the transcriptional changes induced by the oncogene in our system, we performed gene set enrichment analysis (GSEA) comparing the transcriptome of EF-heMSCs to other cell systems that had previously recapitulated the transcriptional signature of the ESFTs to varying degrees. These analysis showed that the EF-heMSC transcriptome is enriched in genes induced by EWS-FLI1 in human bone marrow stromal cells ^22^, and in a rhabdomyosarcoma cell line ^10^(Figure 2G; Table S2). Furthermore, EF-heMSC transcriptome is enriched in genes induced by the oncogene in hpMSCs ^8^, which constitutes thus far the most accepted cellular context of ES ^14^(Figure 2G; Table S2). GSEA using a collection of publicly available soft tissue tumor gene expression profile database ^23^ and genes expressed in ES ^11^ demonstrated that EWS-FLI1 expression in heMSCs induced a transcriptional pattern enriched in genes of the ES signature, but not to other sarcomas like osteosarcoma or rhabdomyosarcoma (Figure 2H; Table S2).

To further characterize the overlap of EF-heMSCs and ESFT transcriptomes, we performed a hierarchical unsupervised analysis, using heSCs and hMSCs jointly as a reference for ESFT transcriptional profiles deposited in GEO (Figure 2I, 2J; Table S3), and selected the 400 most up and down represented genes for comparisons. GSEA revealed the significant enrichment of EF-heMSC transcriptomes in the expression of the highly expressed genes in ESFTs, while genes repressed by the oncogene in EF-heMSCs were among those transcripts underrepresented in ESFTs (Figure 2K; Table S3). Altogether, these results validated the use of the heMSCs as a model to experimentally recreate a *bona fide* ESFT signature.

### EWS-FLI1 binds to Repetitive Genomic Regions in heMSCs

To identify binding sites of EWS-FLI1 in heMSCs, we performed chromatin immunoprecipitation with a Flag antibody to pull down uniquely the Flag-tagged oncogene (avoiding contamination with endogenous FLI1-bound regions), followed by genome wide sequencing (ChIP-seq). This analysis resulted 3086 peaks (2836 annotated peaks) (Table S4). Genome mapping revealed preferential binding of EWS-FLI1 to intergenic and intragenic regions, particularly within introns. Indeed, 1065 peaks in genes corresponded to intronic positions, while 1186 peaks corresponded to intergenic locations (Figure 3A). Characterization of the chromatin states by overlapping the identified peaks and the histone mark profiles in MSCs from the Epigenome Roadmap ^24^ revealed that 45% of the chromatin bound by EWS-FLI1 displayed low levels of all histone marks (quiescent chromatin state), and enrichment of EWS-FLI1 binding to weakly transcribed (TxWk) genomic regions (Figure 3B). Visualization of the distribution relative to the transcription start site (TSS) showed that EWS-FLI1 preferentially binds to regions around 10-100kb of the TSS (Figure 3C). *De novo* motif analysis revealed that the primary DNA sequence underlying EWS-FLI1 binding sites upstream and downstream the TSS is divergent: while the fusion oncoprotein binds to GGAA repeats in intergenic regions, the preferential binding sequence in intron and promoter regions is the tandem iteration of 10 or more CA dinucleotides (Figure 3D).

**Figure 3.**
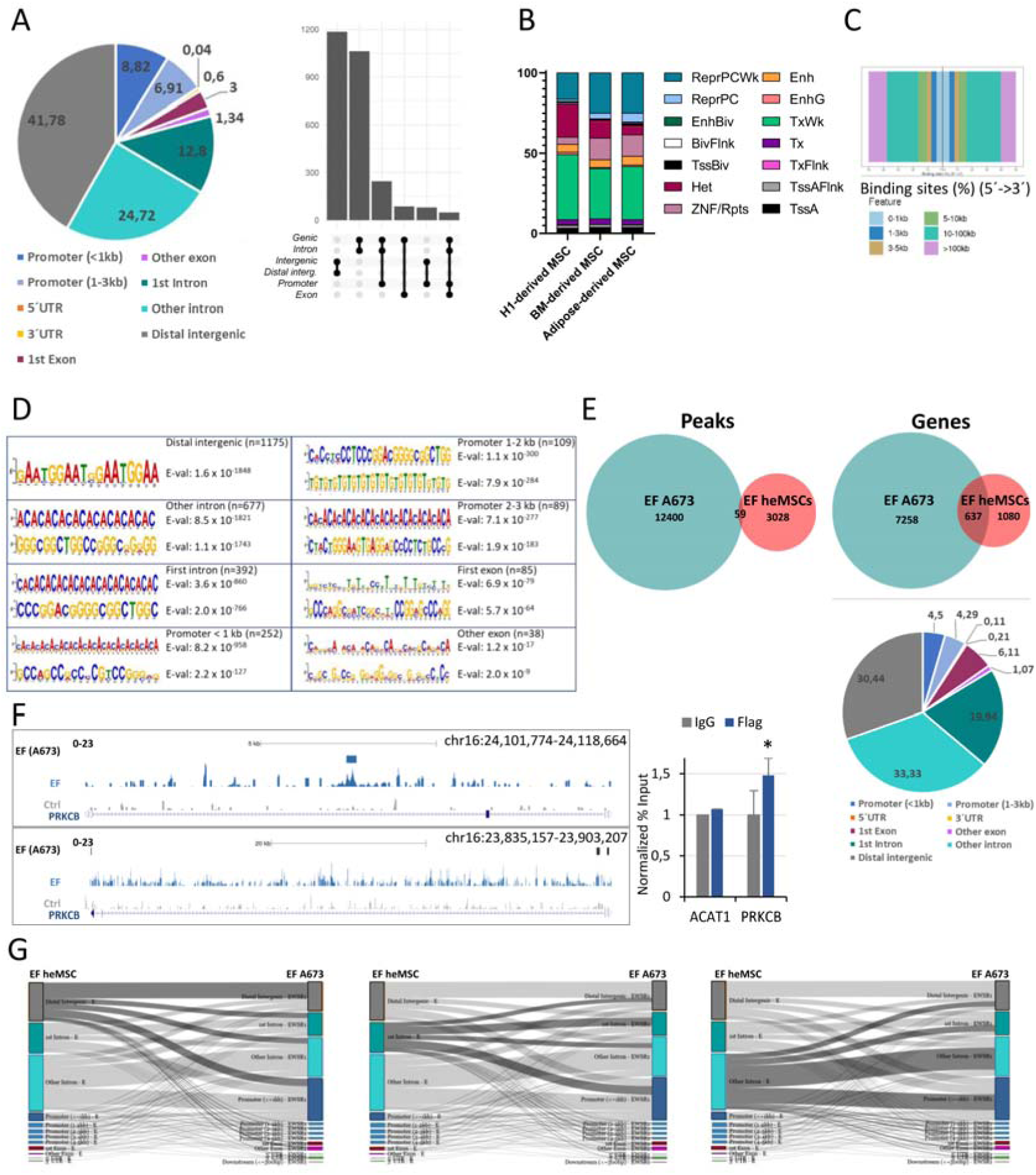
Identification and Characterization of EWS-FLI1 Binding Sites in heMSCs. (A) Left, pie chart depicting genomic annotation of oncogene-bound regions in heMSC-1 cells 48 hours after infection with a Flag-tagged EWS-FLI1, identified by ChIP-seq performed with a Flag antibody. Right, plot for peak annotation overlap. BAM files were merged using BEDTools ^52^, and peak calling using the input as control was performed with MACS2 ^53^. Annotation of the peaks was done using the ChIPseeker ^54^ package. (B) Annotation of chromatin states associated to EWS-FLI1-bound peaks in heMSC-1 cells, performed with BEDtools using chromatin states from the Epigenome Roadmap ^24^ from the following cell lines: E006 (H1 Derived MSCs), E026 (Bone Marrow Derived Cultured MSCs) and E025 (Adipose Derived MSC Cultured Cells). The states are as follows: TssA (Active TSS); TssAFlnk (Flanking Active TSS); TxFlnk (Transcription at gene 5’ and 3’); Tx (Strong transcription); TxWk (Weak transcription); EnhG (Genic enhancers); Enh (Enhancers); ZNF/Rpts (ZNF genes and repeats); Het (Heterochromatin); TssBiv (Bivalent/Poised TSS); BivFlnk (Flanking Bivalent TSS/Enh); EnhBiv (Bivalent Enhancer); ReprPC (Repressed PolyComb); ReprPCWk (Weak Repressed PolyComb). Quies (Quiescent/Low) chromatin state was excluded from this graph for the sake of better visualization of the data. (C) Distribution of EWS-FLI1-binding regions in heMSC-1 cells relative to transcriptional start sites (TSS). (D) Identification of EWS-FLI1 binding motifs by MEME tools ^55^ after classification according to their annotation. (E) Venn diagram illustrating the overlap of genes associated with EWS-FLI1 binding peaks in heMSC and A673 cells. Bottom panel, pie chart illustrating annotations of EWS-FLI1 peaks in heMSCs corresponding to oncogene bound genes. (F) Genome browser tracks depicting EWS-FLI1 binding to the *PRKCB* locus in heMSCs (top screenshot) and A673 cells (bottom screenshot)^25^. Blue tracks, ChIP-seq corresponding to heMSC-1 cells infected with EWS-FLI1; grey tracks, heMSC-1 infected with control lentiviral supernatants. The scale of the tracks (0-23) is the same size for the control and the EF. Right panel, validation of EWS-FLI1 binding to intron 7 of *PRKCB* in heMSCs, detected by ChIP-qPCR in heMSC-1 cells infected with the Flag-tagged oncogene. Values were referred to the percentage of input and normalized with respect to the control condition, and correspond to two independent experiments performed in triplicate. *ACAT1*, negative control. Student’s t-test, *, p<0.05. (G) Sankey plots showing the annotations of peaks corresponding to genes bound by EWS-FLI1 in both heMSC and A673 cells. In each of the plots the transitions from distal intergenic (on the left), first intron (in the middle) and other introns (on the right) of the oncogene peaks in heMSCs to the peaks in A673 cells have been highlighted.

Because EWS-FLI1 induces an ES transcriptome in heMSCs (see Figure 2), we wanted to know whether these transcriptional changes in heMSCs were due to direct EWS-FLI1 binding activity. Strikingly, only few EWS-FLI1-bound genes displayed significant transcriptional changes in heMSCs (Figures S3B, S3C). We tested whether EWS-FLI1 directly binds to the same genomic regions as in a transformed ES cells by overlapping EWS-FLI1 peaks in heMSCs with those peaks identified by other investigators in A673 cells ^25^. Although the overlap between peaks was virtually nonexistent (Table S5), the analysis based on genes demonstrates that 37% of the genes bound by EWS-FLI1 in heMSCs are bound in A673 cells (Figure 3E, Table S5). Importantly, EWS-FLI1 binding in heMSCs occurs in intronic regions in more than 50% of these common genes (Figure 3E). Among these genes is *PRKCB*, which exhibits an EWS-FLI1 binding peak at intron 7 in heMSCs, while in A673 cells the fusion oncoprotein binds to the TSS and the second intron, distant chromatin regions (Figure 3F). To determine the transitions from oncogene peaks in heMSCs to oncogene peaks in tumor cells, we generated Sankey plots that show, for each overlapped gene between samples, which annotations relate to each individual sample (Figure 3G). The illustration of the peak annotation in each sample for each shared gene shows a portion (22%) of genes in which EWS-FLI1 binds in intergenic and intronic regions in heMSCs enriched with the oncoprotein in the 1kb upstream promoter region in A673 cells (Figure 3G). Particularly, this transition occurs between intron peaks in heMSCs and 1kb promoters in tumor cells (16%). These results describe the early steps of EWS-FLI1 reprograming of transcriptome in the cell of origin whereby the binding of the oncogene to intronic regions would promote the formation of chromatin loops leading to ES characteristic GGAA repeats based neo-enhancers.

### EWS-FLI1 Induces BRCA1 Expression and Impairs the DNA Damage Response in heMSCs

ES cells have robust BRCA1 mRNA levels ^26^, which are dependent on EWS-FLI1 expression ^27^. Accordingly, BRCA1 expression detected by immunohistochemistry revealed strong BRCA1 signal in primary ES samples, but not in other developmental tumors, such as rhabdomyosarcoma or neuroblastoma (Figure S4).

Although the number of peaks in exons identified by ChIP-seq in EF-heMSCS is low (4% of the peaks), one of the most extensive EWS-FLI1 binding region was found in large central exon 11 of *BRCA1*. The oncogene also binds to exon 15 of *BRCA1*. These enrichments were confirmed by ChIP-qPCR (Figure 4A). RT-qPCR with different primer pairs to amplify exons 11 and 15 revealed that EF-heMSCs have significantly higher BRCA1 mRNA levels. BRCA1 protein levels were confirmed to be higher in EF-heMSCs (Figure 4B). However, and despite the upregulation of BRCA1 expression, analysis of the DNA damage by the alkaline comet assay revealed that EF-heMSCs harbor defects in DNA damage repair (DDR) mechanisms (Figure 4C).

**Figure 4.**
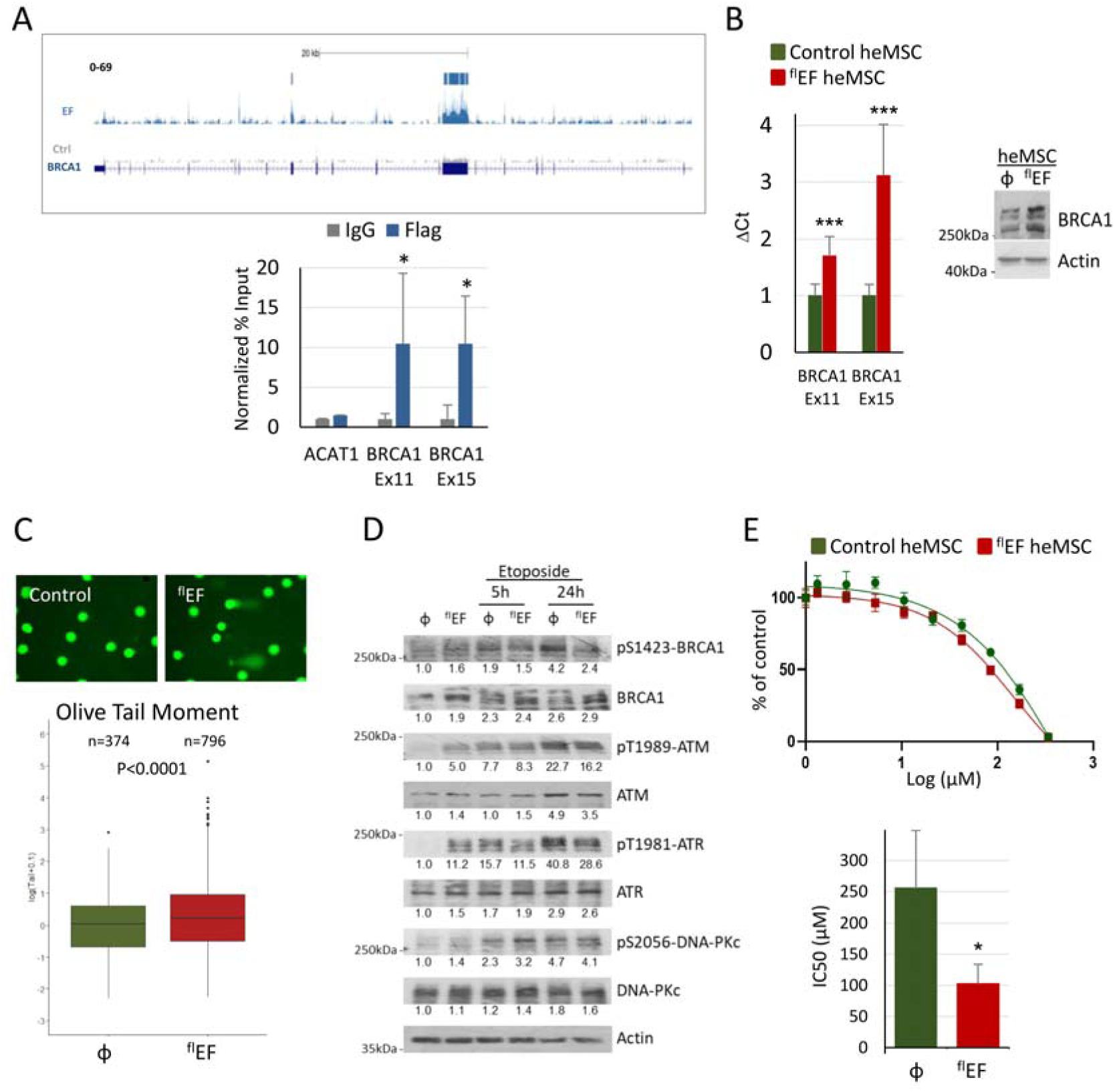
EWS-FLI1 Binds to *BRCA1*. (A) Top panel, genome browser screenshot illustrating EWS-FLI1 binding to the *BRCA1* locus in control and EF-heMSC cells. The scale of the tracks is the same size for the control and the EF Lower panel, chromatin immunoprecipitation of *BRCA1* exons 11 and 15 by EWS-FLI1 in heMSC-1 cells infected with EWS-FLI1. Values were referred to the percentage of input and normalized with respect to the control condition, and correspond to two independent experiments performed in duplicate. ACAT1, negative control. Student’s t-test, *, p<0.05. (B) BRCA1 expression in heMSC-1 cells infected with EWS-FLI1, detected by RT-qPCR and Western blot. Data obtained from three independent experiments performed in triplicate. Student’s t-test, ***, p<0.001. (C) Representative images of the alkaline comet assay performed with heMSC-1 cells infected with control or EWS-FLI1 lentiviral supernatants. Below, Box-Whisker plot representation of the quantification of the product of the tail length and the fraction of total DNA in the tail (Olive tail moment) in control and EF-heMSC cells. Differences between groups were analyzed by using a multiple regression model and a log(x+0.1) transformation. (D) Western blot analysis to detect the expression and phosphorylation status of BRCA1 and kinases involved in DNA damage repair (DDR) in control and EF-heMSC cells under basal conditions and after etoposide treatment. At the bottom, densitometric quantification of Western blot signals, normalized with respect to Actin intensity. (E) Dose-response curves and IC50 values for etoposide in control heMSC-1 and in heMSC-1 expressing EWS-FLI1. Values are representative of three independent experiments. Student’s t-test, * < 0.05.

Since ES cells are highly sensitive to chemotherapeutic agents, we investigated whether defective DDR pathways in EF-heMSCs were related to baseline altered BRCA1 activation or to abnormal response to the treatment with DNA damaging agents like etoposide. EF-heMSC showed upregulation of BRCA1 expression in all experimental conditions. However, BRCA1 phosphorylation in EF-heMSCs was only slightly increased in basal conditions and after etoposide treatment did not reach the phosphorylation levels of control cells (Figure 4D). BRCA1 is the phosphorylation substrate of different kinases, including ATM and ATR, which in turn are activated by phosphorylation at early stages of DNA damage. Expression of EWS-FLI1 in heMSCs induced the phosphorylation of both kinases in basal conditions. However, ATM and ATR phosphorylation was defective in EF-heMSCs upon etoposide treatment (Figure 4D). In contrast, the activation of DNA-PKc, the third kinase, which, along with ATM and ATR, regulates DDR signaling, was unaffected by EWS-FLI1 expression in heMSCs. In support of the notion that EWS-FLI1 expression induces defective DDR, thereby sensitizing tumor cells to the effects of DNA damaging agents, viability assays showed that the IC50 of etoposide is over half-fold lower in EF-heMSCs compared to control heMSCs (Figure 4E).

### EF-heMSCs Form Ewing-like Tumors *In Vivo*

To test the potential *in vivo* tumorigenic effect, heMSCs were collected two days after infection with Flag-tagged EWS-FLI1 and injected into the gastrocnemius of 21-day-old NOD/SCID mice. Few months later (mean latency 5 months), 40% of the mice inoculated with EF-heMSCs showed health problems: they appeared sad and dull, and in some cases pain and breathing difficulties. Post-mortem examination revealed the varying presence of soft, whitish masses in the spine; hematogenous masses in the abdominal cavity affecting the mesothelium, the liver, and other organs; and nodules in the lungs (Figure S5A). Importantly, no such lesions were observed in any of the mice inoculated with control heMSCs. Histologic characterization of an inoculation site revealed the presence of a small nodule with a heterogeneous population of cells among which some aberrant nuclei could be distinguished, while the lumps in the posterior aspect of the spine were composed of tubular structures diffusely infiltrating the fat formed by immature lipoblasts which in the periphery of the tumor masses show a myxoid appearance (Figure 5A). Of note, the nuclei of the cells in the vessels were round to oval and showed a distinctive “salt and pepper” chromatin. The masses in the abdominal cavity showed characteristic features of ESFT: densely packed, small, uniform and poorly differentiated cells with scant cytoplasm and round nuclei with frequent areas of hemorrhage, necrosis, and scattered mitotic figures. In addition, in one of the animals, tumor cells were observed to massively grow in the spine (Figure 5A).

**Figure 5.**
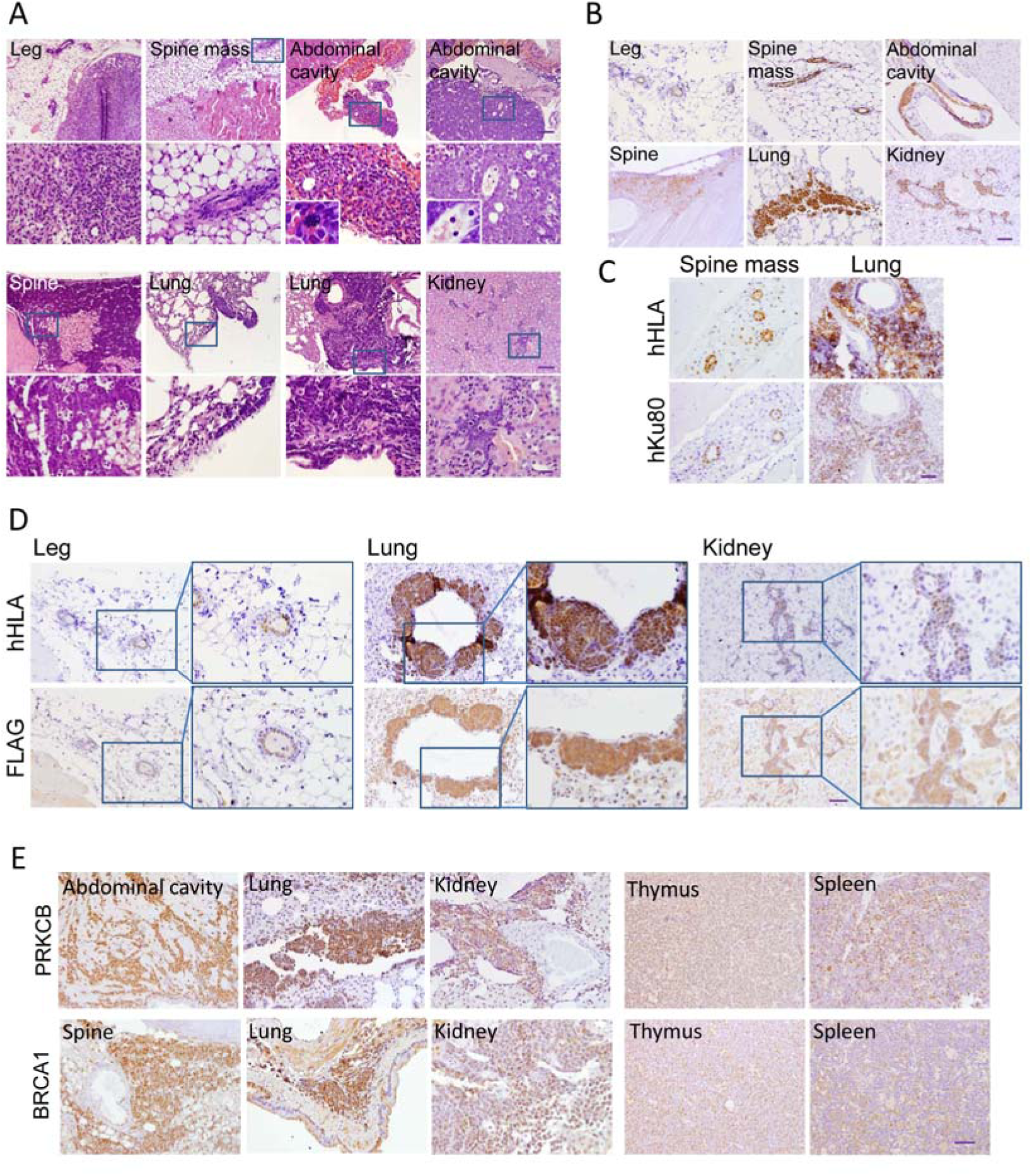
EF-heMSCs Form Ewing-Type Sarcomas In Vivo. (A) H&E staining of representative lesions from NOD/SCID mice injected in the gastrocnemius with EF-heMSCs. Images show representative sections at two different magnifications (top bars, 100µm; bottom bars, 20µm). Observe the aberrant mitotic figure and the presence of blood lakes shown in the insets of the abdominal tumor masses. (B) IHC to detect human HLA. Magnification bars: 50μm. (C) IHC to detect human HLA and human Ku80 in sequential sections of the same sample. Magnification bars: 50μm. (D) IHC of human HLA and Flag in sequential sections of EF-heMSC samples. (E) IHC of PRKCB and BRCA1 in EF-heMSC-derived tumors. Thymus and spleen sections correspond to normal tissues. Magnification bars: 50μm.

Histologic examination of the different organs revealed metastatic dissemination to the lungs, where the cells initially colonized the pleura and spread to the bronchial tree and bronchioles, and to the liver and kidney parenchymás (Figure 5A). Positive human HLA staining was observed in cells of the vascular-like tubes, and in cells from primary tumors and from pulmonary and renal metastases (Figure 5B). Subsequent sections subjected to immunohistochemical staining for human HLA and human Ku80 further confirmed the human nature of these cells (Figure 5C and Figure S5B).

To determine EWS-FLI1 expression in these lesions, sections were immunostained with the Flag antibody. EWS-FLI1 expression could be detected in the vascular formations and in the neoplastic lesions (Figures S5C, S5D and S5E). Immunohistochemical staining of adjacent sections with human HLA or Flag antibodies corroborated that the Flag signal corresponded to human HLA-positive cells (Figure 5D). In addition, we confirmed the expression of PRKCB and BRCA1, direct targets of EWS-FLI1 in heMSCs, in these lesions (Figure 5E). Recapitulating, vascular structures and primary and metastatic neoplastic lesions were formed by human cells expressing the oncogene.

### The Transcriptional Profile of the Experimental Ewing Tumors Resembles Ewing Sarcoma

To characterize the transcriptional signature of the experimental tumors we performed single-cell RNA sequencing on formalin-fixed tumors using the 10X Genomics Visium platform, which allows also for the reconstruction of tissue organization. Unsupervised clustering of gene expression led to the identification of tumor cells, as shown in the spatial DimPlots of Figure 6A. Spatial maps of the expression of some of the most relevant genes by uniform manifold approximation and projection (UMAP) dimensional reduction confirmed the expression of those genes in tumor cells (Figure 6B). Interestingly, the expression of these genes in ES cell lines was higher than in any other tumor cell line tested (Figure 6C), many already described in A673 cells as direct targets of the oncogene ^25^.

**Figure 6.**
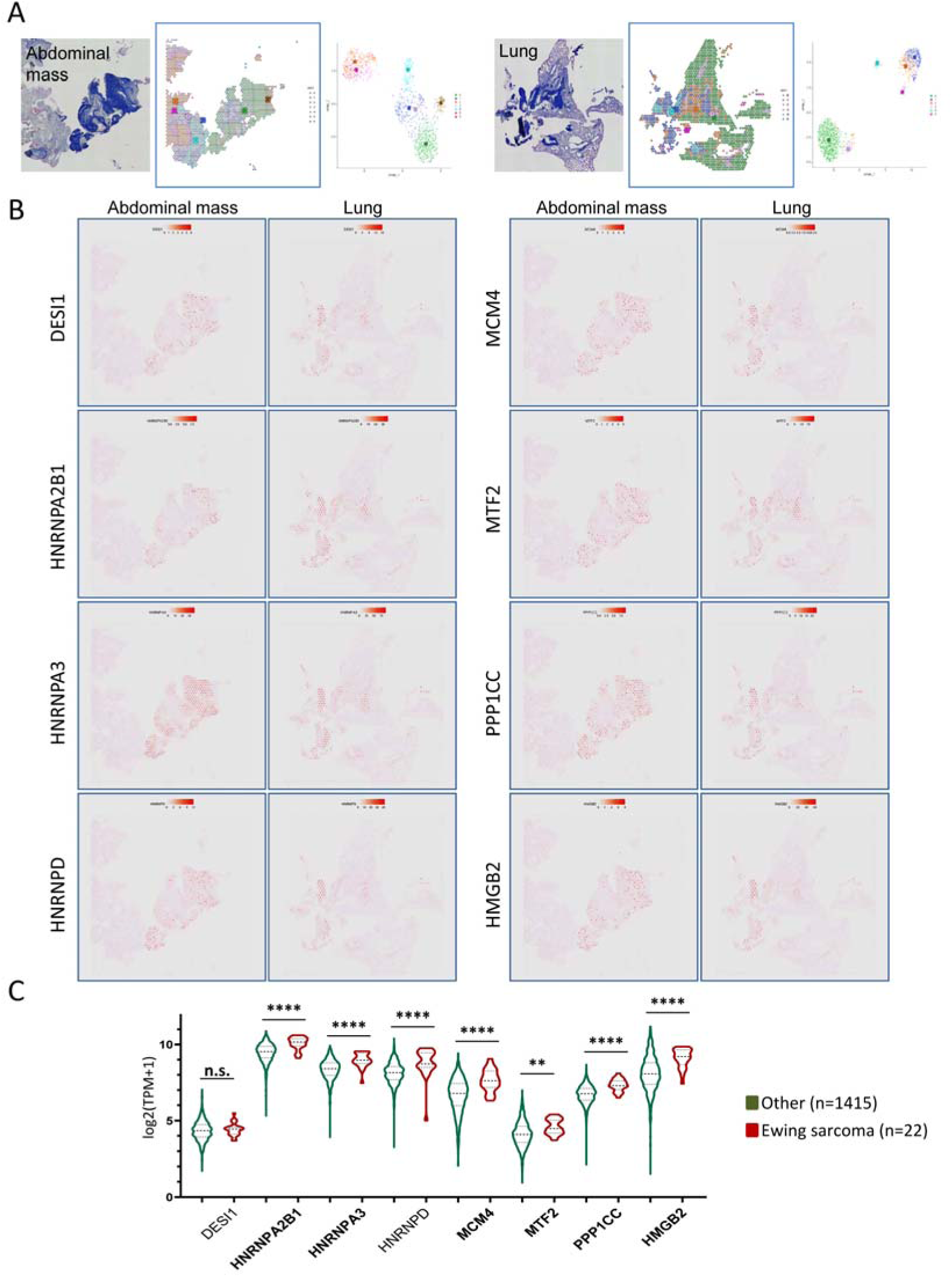
Transcriptional Characterization of Experimental Ewing Sarcoma Tumors. (A) H&E staining of an abdominal mass and a lung metastasis, and identification of cell clusters based on their corresponding UMAPs. (B) Expression maps of some of the most relevant genes that determine clusterization in the above UMAPs. (C) Expression levels in Ewing sarcoma cell lines and in other tumor cell lines, extracted from DepMap portal (https://depmap.org/), of the most relevant genes identified by computation of cell clusters from UMAP dimensional reduction. Legends in bold highlight genes directly bound by EWS-FLI1 in A673 cells in promoters or enhancers ^25^.

With the aim of finding markers specific for ESFT, we took advantage of the unsupervised analysis using heSCs and hMSCs to distinguish the ESFT transcriptional profiles (see above, Figure 2G and Table S3). The top 100 DEGs (adjusted p-value < 0.05) considering the log2(FC) were filtered, and spatial expression images generated for those genes that were cross-represented in both datafiles. As shown in Figure 7A, both the primary tumor and the lung metastasis shared the expression of many of those genes. Among them, BCL11B and ITM2A, previously identified as genes specifically related to neural and endothelial features of ESFTs ^11^; expressed in most ESFT specimens; and undetectable or weakly expressed in other developmental tumors (Figure S5F). Importantly, BCL11B and ITM2A were immunodetected in the experimental tumors, whereas staining of normal tissues such as thymus, spleen or liver resulted in no or low expression (Figure 7B).

**Figure 7.**
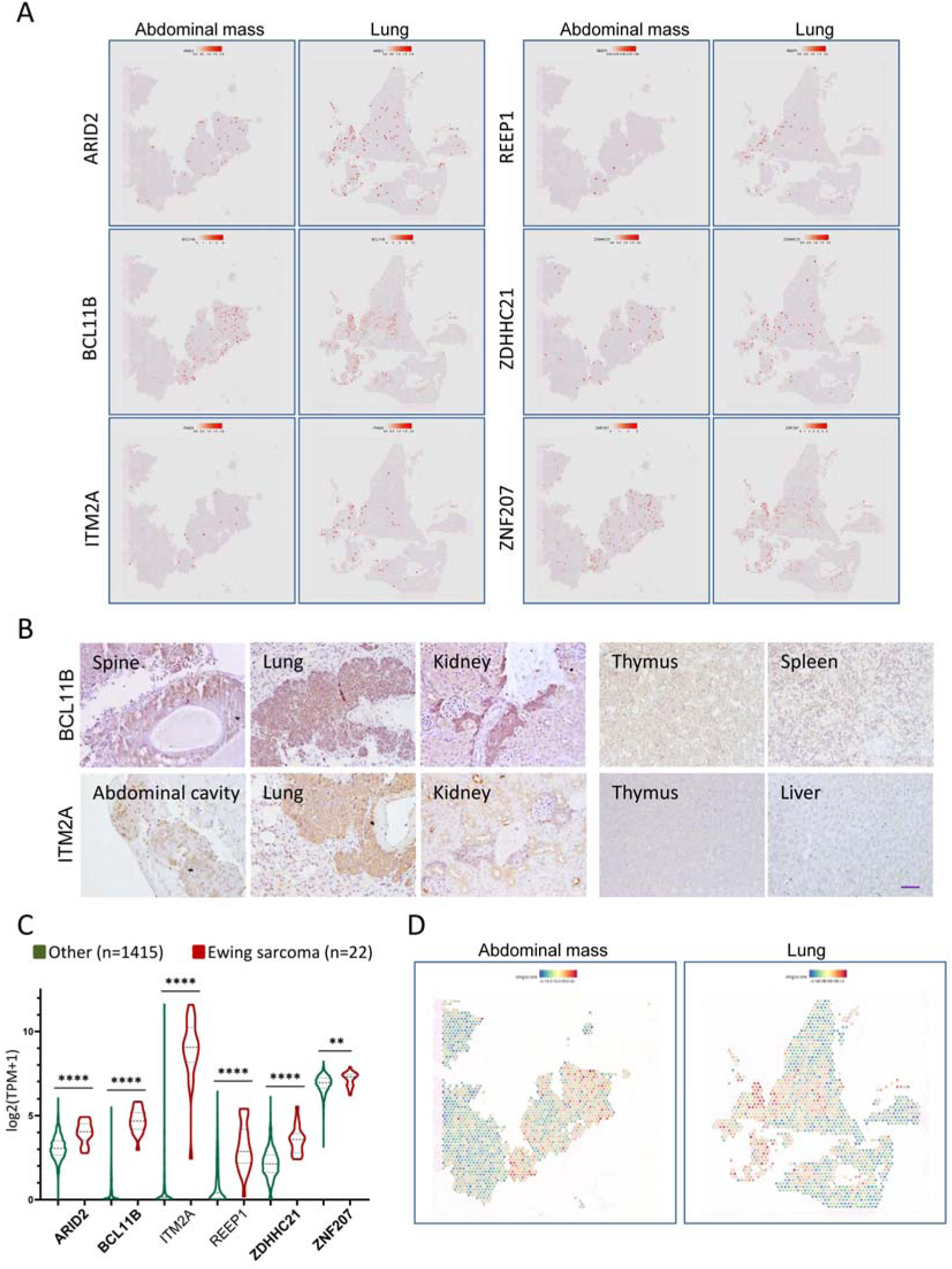
ESFT Transcriptional Signature of EF-heMSC Derived Experimental Tumors. (A) Expression maps of some of the genes differentially expressed in Ewing sarcomas (Table S3) (adjusted p-value <0.05, log2(FC), top 100). (B) IHC of the ESFT distinctive markers BCL11B and ITM2A in EF-heMSC-derived tumors. Thymus, spleen and liver sections correspond to normal tissues. Magnification bar, 50μm. (C) Expression levels of differentially expressed genes in Ewing sarcoma cell lines and in other tumor cell lines, extracted from DepMap portal (https://depmap.org/). Legends in bold highlight genes directly bound by EWS-FLI1 in A673 cells in promoters or enhancers ^25^. (D) Spatial images of the gene set scores obtained by the singscore package ^57,58^. As reference gene set, the top 400 (for abdominal mass) or 200 (for pulmonary metastasis) differentially expressed genes in Ewing sarcoma (Table S3) were ordered by the log2(FC) (adjusted p-value <0.05).

To further assess the specificity of the identified ESFT markers, we investigated their expression in cancer cell lines. DepMap data analysis revealed that they were all significantly overexpressed in ES cell lines compared to other tumor cell lines (Figure 7C). Finally, spatial DimPlots representing the gene signature score for each cell using the most differentially expressed ESFT genes (Table S3; adjusted p-value <0.05; log2(FC)) as reference, confirmed the transcriptional similarity of the experimental tumors to human ES (Figure 7D).

## DISCUSSION

ESFTs are highly undifferentiated developmental cancers with a translocation as the only pathognomonic genetic feature (for a recent review, see ^28^). The oncoprotein resulting from this translocation is the tumor cell driver of tumorigenesis responsible for blocking cell differentiation and inducing cell transformation. Our working hypothesis was that the sole expression of the oncogene should be sufficient to induce an ES-specific transcriptional signature and generate tumors when expressed in the rightly undifferentiated embryonic stem-cell context, such as a progenitor in the stage of mesenchymal and/or endothelial differentiation during fetal transitions at gastrulation ^12^. Indeed, mesenchymal cells derived from the mesoderm or the neural crest are currently considered the most likely candidates as cells of origin of ESFTs ^28,29^. Previous work by Riggi *et al.* using hMSCs showed that the degree of differentiation, or in this case, undifferentiation, of the cells is relevant for oncogene-induced transcriptional reprogramming ^14^. Our experimental approach consisted of using human embryonic mesenchymal cells (heMSCs) from experimental teratomas. The analysis of these progenitor cells clearly showed how heMSCs have multilineage potentials and thus are less differentiated compared to hpMSCs, already constrained towards bone and chondrogenic lineages.

Low levels of the Flag-tagged oncogene in heMSCs were sufficient to profoundly alter the transcriptional profile of these cells. In addition to inducing or repressing previously recognized target genes, EWS-FLI1 induced the transcription of numerous vascular differentiation genes and, to a lesser extent, of genes specific to neural lineage. This EWS-FLI1-induced transcriptional signature in heMSCs recreated the transcriptome of ES, implying that the oncogene is able to impose in the cell of origin an aberrant hybrid differentiation program, with endothelial and neural characteristics. Our results confirm previous findings demonstrating that the ESFT-specific expression profile is enriched in endothelial and neural genes ^11^. Therefore, the ESFT transcriptome is a direct consequence of the activity of the oncogene on cell differentiation, and does not reflect the histogenesis of these tumors. Strikingly, despite inducing a transcriptional signature characteristic of ES, the chromatin binding pattern of EWS-FLI1 in heMSCs is distinct from that in the ES cell line A673. In primary hMSCs, sites activated by EWS-FLI1 have a closed chromatin conformation that switch to an open chromatin conformation upon EWS-FLI1 expression, which acts as a pioneer factor. The recruitment of the transcriptional machinery results in an active enhancer pattern analogous to the chromatin architecture of ES ^30^. Our results suggest that the ability of EWS-FLI1 to enhance chromatin accessibility in the cell of origin, i.e., the heMSC, may be initially triggered by its binding to CA microsatellites in intronic regions distinct from GGAA multimers, permitting the establishment of long-range interactions which eventually will cause an EWS-FLI1 enrichment in GGAA repeat elements at enhancers in ES cells. In this regard, it has been shown that the transcription factor NCoA3 governs the dynamic chromatin landscape through its binding to internal body sequences, probably by keeping the enhancer and promoter in close proximity to the intronic sequences recognized by NCoA3 ^31^. Since the frequency of microsatellites is not different in distinct chromatin regions, except for those with CG repeats which are sensitive to methylation ^32^, the question arises as to which factor or factors determine the binding affinity of EWS-FLI1 to specific DNA sequences depending on the intergenic or intronic locations. In any case, our results support the notion that the neo-enhancer looping is a critical process in ES pathogenesis ^30,33^.

Regulatory GGAA microsatellite repeats are essential for maintaining the oncogenic properties of ES cells ^30^. However, it is unlikely that EWS-FLI1 exclusively uses this mechanism to unleash the plethora of cellular changes that will ultimately result in ES. Moreover, only a fraction of the direct target genes of EWS-FLI1 have been identified so far, mostly in ES cell lines, which limits our understanding of the mechanisms of action of the fusion oncoprotein and its impact in early stages of tumorigenesis ^33^. A well-known but little understood feature of ESFTs is their intrinsic chemo- and radio sensitivity, at least initially. Some authors have proposed ESFTs as BRCA1-deficient tumors since EWS-FLI1 increases transcription to cause R-loops and block BRCA1 repair ^26^. In contrast, other studies have shown the survival dependence of ES cells on BRCA1 expression and claim for a functional ATM deficiency as a major DNA repair defect and ATR as a collateral dependency ^27^. Appreciating the considerable differences between these studies, all based exclusively on cell lines exclusively with a broad range of different mutations that may affect DDR, our results would support the latter hypothesis, since we demonstrate that BRCA1 is a direct target of EWS-FLI1 in the potential cell of origin heMSCs. Consistent with this result, heMSCs expressing the oncogene have elevated levels of BRCA1 and are concomitantly deficient in DDR pathways, including defects in BRCA1 phosphorylation by ATM and ATR. Our working hypothesis is that the oncogene could be inducing the expression of phosphatases whose hyperactivity might decrease the ATM/ATR phosphorylation levels. Regardless of the underlying mechanism, these results demonstrate that the mere presence of the oncogene is responsible for the intrinsic vulnerability of ES cells to DNA damaging agents.

Our efforts to maintain EWS-FLI1-infected heMSCs in culture have been unsuccessful, likely due to the rapid induction of p53-p21-RB1 signaling. It has been reported that EWS-FLI1 induces p53-mediated growth arrest in human primary fibroblasts ^34^, and the growth arrest can be attenuated in p16 or p53 deficient mouse embryonic fibroblasts ^35^. However, the p53 pathway and DNA damage signaling pathway are functionally intact in ESFT ^36^, and to date, the only cells in which EWS-FLI1 does not induce cell cycle arrest and can be stably expressed are stem-like progenitor cells ^7^. In our model, the discordance between the anti-proliferative effects in our model and the tolerance of EWS-FLI1 expression could be related to the oncogene concomitantly regulating the expression of proliferation and survival genes critical for heMSC viability. Alternatively, this discrepancy could be attributable to a delicate balance of expression between pro-survival and cycle arrest genes dependent on the levels of oncogene expression.

*In vivo*, EWS-FLI1-expressing heMSCs generated frequently hemorrhagic bone and soft tissue tumors, in accordance with the observation that ESFTs arising in soft tissues are morphologically and molecularly indistinguishable from those arising in bones ^37^. These results indicate that the same cell of origin would be capable of giving rise to both clinical presentations. In addition to the expression of markers previously described as molecular discriminants of ESFTs, our experimental tumors display a histology compatible with human ESFTs. Collectively, all these findings would endorse James Ewing’s characterization, one hundred years ago, of this group of sarcomas as diffuse endotheliomas of bone and explain his original hypothesis of an endothelial cell of origin ^4^.

In this study we have isolated and characterized human embryonic mesenchymal cells that fulfil the criteria for the cell of origin of ES. Our results suggest that EWS-FLI1 is primarily a transcription factor with a marked potential to induce an aberrant cell differentiation program during early embryonic development with limited tumorigenic capacity. We hypothesize that changes in oncogene expression levels during puberty, potentially through hormonal influence, increase the chances for ES tumors to develop. This would explain our previous observation of *EWSR1* rearrangements in hemangioma samples from prepubertal patients who subsequently developed ES during pubertal growth ^38^.

In conclusion, we propose early human embryonic mesenchymal stem cells as the cell of origin of ES. These cells present a distinct binding pattern of the oncogene with intronic microsatellites (>10 CA dinucleotides) as the preferential sites; show defects in DDR and increased expression of BRCA1; and form tumors in mice with characteristics of human ES, i.e. small, round, undifferentiated cells expressing characteristic ES markers. Our experimental model reproduces the clinicopathological characteristics of ES and allows for the characterization of early tumorigenesis mechanisms and the discovery of new markers of disease.

## METHODS

### Cell Culture

Commercial cell lines were grown in standard conditions. The heSC line ES4 was previously characterized, and maintained as described ^39^. heMSCs, isolated from teratomas by enzymatic digestion, were grown in Eagle’s medium supplemented with 10% FBS and bFGF (1ng/ml). ES4 and heMSC characterization was performed in November 2018 (qGenomics, Spain). A673 and 293T cells were cultured in standard conditions. Pediatric hMSCs were obtained from bone marrow aspirates of HSJD pediatric normal subjects under written informed consent.

### Teratoma Formation

Experimental teratomas in NOD/SCID mice (Charles River) were generated as previously described ^40^. Animal experiments were conducted following protocols approved by the Institutional Ethics Committee on Experimental Animals, in full compliance with Spanish and European laws and regulations. Immunofluorescence characterization of the experimental teratomas was performed with antibodies to detect TuJ1 (Covance), FOXA2 (R&D Systems), AFP (Dako), and α-SMA, α-sarcomeric actin and NF-200 (Sigma).

### heMSC Phenotyping and Trilineage Differentiation

Immunophenotypic analysis was carried out using antibodies to detect CD31 (Miltenyi Biotec), and CD73, CD90, CD34, CD45, CD105 and HLA-I (BD Biosciences) with a LSRII flow cytometer running FACSDiva v5.02 Software (BD Biosciences). Trilineage differentiation was performed with the StemMACS™ Trilineage Differentiation Kit human (Miltenyi Biotech), as described before ^41^.

### *In Vitro* Functional Assays

To assess *in vitro* growth rates, 1×10^5^ cells per 10-cm plate were seeded. Every two or three days, cells were harvested, and 1×10^5^ cells were replated until cells stopped proliferating. The population doubling time was calculated as described ^42^.

### RT-qPCR

RNA was isolated with Genelute Total Mammalian RNA Kit (Sigma-Aldrich) and cDNA was obtained by using Transcriptor First Strand cDNA Synthesis Kit (Roche). RT-qPCR assays were performed using SYBR Green PCR master mix (Applied Biosystems, Life Technologies). For normalization purposes, we run simultaneously RT-qPCR with primers for GAPDH. The ABI PRISM 7900HT cycler’s software calculated a threshold cycle number (Ct) at which each PCR amplification reached a significant threshold level.

Primer sequences used were:

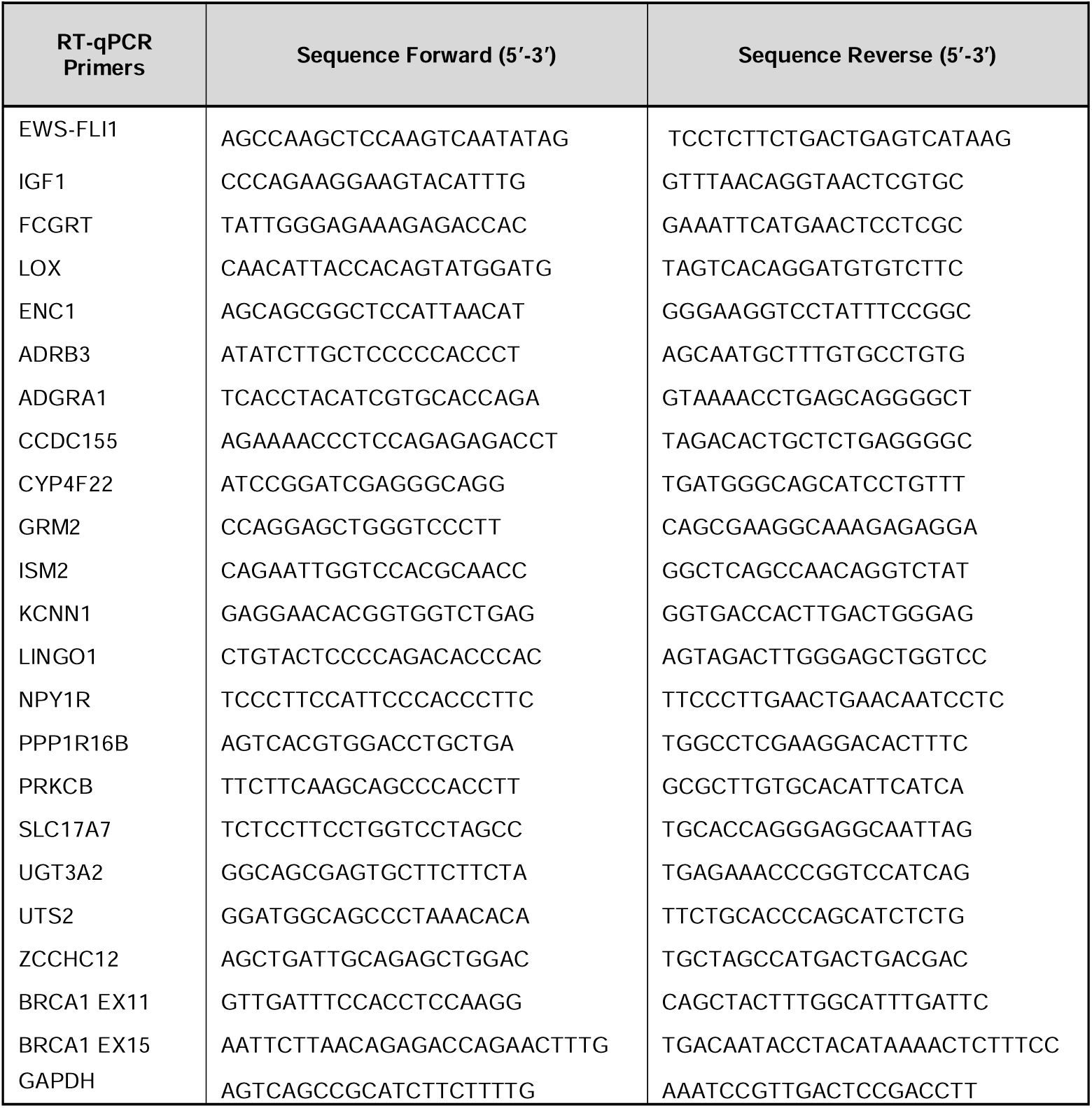

### Immunohistochemistry, Immunofluorescence and Western Blot

These techniques were performed using standard methods. Antibodies used for these techniques were: FLI1, p53, p21, ATM, hHLA, PRKCB (Santa Cruz); CD99, hKu80 (Cell Signaling); β-actin (Abcam); RB1 (BD Biosciences); human nucleus, BRCA1, pS1981ATM (Millipore); pS1423-BRCA1, ATR, pT1989-ATR, DNA-PKc, pS2056-DNA-PKcs (Abclonal); Flag (Merck Life Science); BCL11B (Biolegend) and ITM2A (Fisher Scientific). Approval to conduct this study involving human samples was obtained from the ethic committee from Parc de Salut MAR in accordance with the guidelines of the Helsinki Declaration of 1975, as revised in 1983.

### RNA Sequencing

Total RNA from heMSCs and hpMSC was purified and reverse transcribed. NGS libraries with polyA capture were prepared according to the manual Protocol for use with NEBNext Ultra II Directional RNA Library Prep Kit for Illumina (New England Biolabs) (version 1.0, 7/2 Illumina TruSeq library preparation (Illumina). Libraries were sequenced on an Illumina HiSeq 2500. Raw sequencing reads in the fastq files were mapped with STAR version 2.7.1a ^43^ Gencode release 36 based on the GRCh38.p13 reference genome and the corresponding GTF file. The table of counts was obtained with FeatureCounts function in the package subread, version 1.6.4 ^44^. The differential gene expression analysis (DEG) was assessed with voom+limma in the limma package version 3.48.0 ^45^ using R version 4.1.0. Genes having less than 10 counts in at least 2 samples were excluded from the analysis. Raw library size differences between samples were treated with the weighted “trimmed mean method” TMM ^46^ implemented in the edgeR package ^47^. The normalized counts were used in order to make unsupervised analysis, PCA and clusters. For the differential expression (DE) analysis, read counts were converted to log2-counts-per-million (logCPM) and the mean-variance relationship was modelled with precision weights using voom approach in limma package.

Pre-Ranked Gene Set Enrichment Analysis ^48^ implemented in clusterProfiler ^49^ package version 4.0.0 was used in order to retrieve enriched functional pathways. The ranked list of genes was generated using the - log(p.val)*signFC for each gene from the statistics obtained in the DE analysis with limma ^45^. Functional annotation was obtained based on the enrichment of gene sets belonging to gene set collections in Molecular Signatures Database (MSigDB). The collection used in this project is c5.bp: Gene sets derived from the Biological Process Gene Ontology (GO), version 7.2. RNA-seq data were deposited in GEO under accession number GSE215008.

### Gene Expression Analyses

Raw Affymetrix CEL files were normalized and processed by RMA using the R/Bioconductor oligo package and Affymetrix ^50^. Unsupervised analyses were performed by PCA and hierarchical clustering using Euclidean metrics. Standard deviation (SD) density plots were used to determine the cut-off value for gene expression unsupervised analyses (SD≥1). Supervised gene expression analysis was performed using limma ^45^. Probes were considered significant differentially expressed when the FDR-adjusted Wilcoxon rank test for independent samples was <0.05 and the absolute logarithmic fold change (|LFC|) was >2.

### Gene Set Enrichment Analysis (GSEA)

Functional annotation was obtained based on: 1) Gene sets from collections in Molecular Signatures Database (MSigDB) related to Ewing sarcoma; 2) Gene lists provided by Baird’s et al. ^23^; 3) The TOP 400 up and TOP 400 down-regulated genes derived from public data from GEO datasets GSE6460, GSE7637, GSE7896, GSE8884, GSE9451, GSE9440, GSE9510, GSE9520, GSE9593, GSE10315, GSE13604, GSE13828, GSE17679, GSE34620, GSE37371 and GSE31215 (n=236). Expression was normalized using Quantile Normalization. For the differential expression analysis to obtain the list of interested genes, an empirical Bayes moderated t-statistics model (limma) was built. GSE id variable was included in the model as a covariate. To identify the most variable probes, we selected the probes above 1 standard deviation, which were 8176. Top 400 up and top 400 down-regulated genes were obtained ranking the gene by p.adjusted and logFC values. Gene sets description: MIYAGAWA_TARGETS_OF_EWSR1_ETS_FUSIONS_UP MSigDB; ZHANG_TARGETS_OF_EWSR1_FLI1_FUSION MSigDB; RIGGI_EWING_SARCOMA_PROGENITOR_UP MSigDB; STAEGE_EWING_FAMILY_TUMOR MSigDB; (EWS Baird et al); heSC-vs-MSC_DN Gene sets derived from GEO datasets (Top 400 down genes); heSC-vs-MSC_UP Gene sets derived from GEO datasets (Top 400 up genes).

### ChIP-qPCR, ChIP-seq and Bioinformatic Analysis

Cells were subjected to ChIP-qPCR following standard procedures. Briefly, cells were fixed with crosslink solution (HEPES 50mM pH 8.0; NaCl 100mM; EDTA 1mM; EGTA 0.5mM; 0.5% Formaldehyde) for 10 min at room temperature, and incubated with stop solution (10% Glycine in Tris-HCl 10mM pH 8.0). Cells were lysed for 20 min on ice with 10 mM Tris-HCl pH 8.0, 0.25% Triton X-100, 10 mM EDTA, 0.5 mM EGTA, 20 mM β-glycerol-phosphate, 100 mM NA-orthovanadate, 10 mM Na Butyrate and complete protease inhibitor cocktail. The supernatants were sonicated, centrifuged at 13,000 rpm for 15 min, and supernatants were incubated overnight with the Flag antibody (Merck Life Science) or with anti-mouse IgGs (Millipore).

Precipitates were captured with protein A/G-Sepharose, extensively washed, and the crosslink was reverted by incubating the samples with Proteinase K. DNA was extracted and analyzed by qPCR by using the ABI 7700 sequence detection system and SYBR Green master mix protocol (Applied Biosystems). Each immunoprecipitation was done in triplicate. The reported data represent real-time qPCR values normalized to input DNA and are expressed as percentage (%) of bound/input signal. Primers used for ChIP-qPCR were:

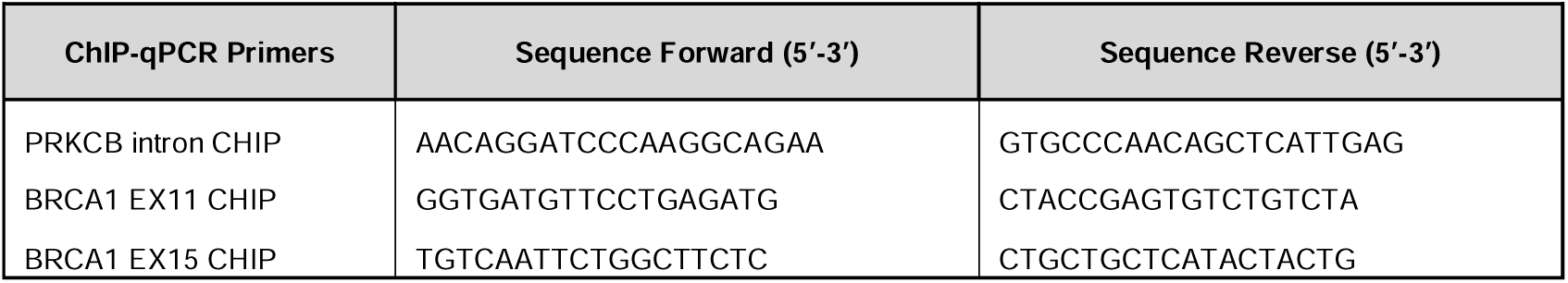

ChIP-seq libraries were prepared using the NEBNext Ultra DNA Library Prep from Illumina according to the manufacturer’s protocol. Briefly, 5 ng of input and ChIP-enriched DNA were subjected to end repair and addition of “A” bases to 3′ ends, ligation of adapters, and USER excision. All purification steps were performed using AgenCourt AMPure XP beads (Qiagen). Library amplification was performed by PCR using NEBNext Multiplex Oligos from Illumina.

Raw sequencing reads in the fastq files were mapped with Bowtie2 ^51^ version 2.4.2 on the GRCh38.p13 reference genome and the corresponding GTF file. BAM files were merged using BEDTools ^52^ version 2.30.0 and the standard parameters, for the different replicates in order to get one file per sample condition. Peak calling was obtained using MACS2 ^53^ tool without building the shifting model (–nomodel flag) and specifying the size of the binding region (–extsize flag) using information from QC (phantompeakqualtools). Annotation of the peaks was done using the ChIPseeker ^54^ package version 3.30.1. GENCODE version 41 was used to annotate the peaks using version 3.16.0 from TxDb.Hsapiens.UCSC.hg38.knownGene package. Ensemble version 86 was used to transform from entrez to symbol annotation of the genes. Annotation of chromatin states was done with BEDtools using the chromatin states from the [Epigenome Roadmap]^24^. We used the files [EXXX]_15_coreMarks_hg38lift_mnemonics.bed.gz from https://egg2.wustl.edu/roadmap/data/byFileType/chromhmmSegmentations/ChmmModels/coreMarks/jointModel/final/.

Overlapping peaks between EWS-FLI1 in heMSCs and FLI1 in A673 ^25^ were identified with the function findOverlapsOfPeaks of the ChIPpeakAnno package (version 3.30.1) using a window of 100bp. Annotation of the peaks was done using the ChIPseeker ^54^ package version 3.30.1. GENCODE version 39 was used to annotate the peaks using version 3.15.0 from TxDb.Hsapiens.UCSC.hg38.knownGene package.

For the identification of consensus binding sites, the peaks were split according to their annotation and fastq files were written using the packages GenomicRanges (version 1.50.2), BSgenome.Hsapiens.UCSC.hg38 (version 1.4.5) and rtracklayer (version 1.58.0). MEMECHIP was then run over the generated fastq files using MEME (version 5.1.1)^55^.

EWS-FLI1 peaks in A673 ^25^ were annotated using annotatePeak function from ChIPseeker (version 1.34.1)^54^ considering TSS range from -5000bp to 100bp and using the annotation from TxDb.Hsapiens.UCSC.hg38.knownGene package (version 3.15.0)(Team, Bioconductor Core, and Bioconductor Package Maintainer. 2021. TxDb.hsapiens.UCSC.hg38.knownGene: Annotation Package for TxDb Object(s). Peak annotation of EWS-FLI1-bound genes shared between heMSCs and A673 were retrieved and analyzed.

All analyses were performed under R version 4.2.1 (R Core Team 2022; https://www.R-project.org/).

### Comet Assay

Cells were infected with lentiviral supernatants and recovered 72h later. Alkaline comet assay was performed using reagents from Trevigen in accordance with the manufacturer’s protocol. Imaging was performed with a fluorescence microscope and Tail Moment determined using the ImageJ analysis software. Fifty or more comets were analyzed for each experiment.

### MTS

Cells were seeded in 96-well plates (1500 cells/well) and infected with lentiviral supernatants. 24h later, cells were treated with serial dilutions of etoposide. Seventy-two hours later, 10% MTS (Promega) was added and absorbance at 490nm was read using a Tecan microplate reader. Percent viability was calculated by normalizing absorbance values to those from cells grown in media with vehicle treatment, after background subtraction. IC50 was determined using Prism 8 Software (GraphPad).

### *In Vivo* Tumorigenesis Assay

heMSCs were infected with Flag-tagged EWS-FLI1 lentiviral supernatants and 48 hours later were recovered. 1×10^6^ cells were imbedded in matrigel and injected into the gastrocnemius of 21 days old NOD/SCID mice (Charles River). Mice were sacrificed when the animals showed clear signs of distress or the presence of any tumoral growth. Dissected tumors and organs were fixed in 10% formalin, and processed routinely for histologic examination.

### Spatial Transcriptomics

Tumor samples were formalin fixed and paraffin embedded. Spatial transcriptomics was performed with Visium CytAssist Spatial Gene Expression Reagent Kits, following the protocol CG000495. RNA quality of the tissue was tested by calculating the percentage of total RNA fragments >200 nucleotides (DV200) of RNA extracted from tissue sections. Once evaluated RNA quality, 5μm tissue sections were placed on SuperFrost slides and placed in a desiccator to ensure proper drying. After overnight drying, samples were deparaffinized, stained with H&E, and images of the tissues were taken, following the protocol Visium CytAssist Spatial Gene Expression for FFPE Deparaffinization, H&E Staining, Imaging and Decrosslinking (CG000520-Rev C). Then, the tissues were decrosslinked and the libraries were constructed following the instructions in the protocol for Visium CytAssist Spatial Gene Expression for FFPE, Human Transcriptome, 6.5mm (Preparation Guide CG000518-Rev D), and sequenced (slide 6,5mm 250MR: 5000 spots at 50000 reads/spot, 40Gb/sample). To analyze the data the spaceranger count function within the 10x Genomics Space Ranger 3.0.0 tool, utilizing the GRCh38-2020 reference genome, was employed. The resulting “filtered_feature_bc_matrix.h5” files were then imported for subsequent analysis, leveraging the Seurat package (version 5.0.3)^56^. The SCTransform function with default parameters was applied to normalize the data, identify variable features, and scale each sample independently. Additionally, PCA dimensionality reduction, determination of k.param nearest neighbors, and computation of cell clusters preceded the implementation of UMAP dimensional reduction. Gene set scores were obtained using the singscore package version 1.18.0 ^57,58^. Data from the transcriptome analysis of human Ewing sarcoma versus stem cells was used to create a gene reference; specifically, the top 100, 200 or 400 differentially expressed genes (adjusted p-value < 0.05) were ordered by the log2(FC). The scaled data from the Seurat object was ranked using rankGenes function. The output was then used for the simpleScore function, determining the direction as known (knownDirection = TRUE). The obtained total score was then imputed to the Seurat object. All analyses were performed with R version 4.2.1 (R Core Team 2022; https://www.R-project.org/).

## DATA AVAILABILITY

Data from array analyses have been deposited in GEO, accession GSE GSE272960 (GSE272957 for RNA-seq, GSE272958 for spatial transcriptomics and GSE272959 for ChIP-seq). To review GEO accession GSE272960, go to https://www.ncbi.nlm.nih.gov/geo/query/acc.cgi?acc=GSE272960.

## ACKNOWLEDGMENTS

The authors would like to thank Toni Ventura, Elisenda Alari-Pahissa, and Isadora Lemos for their technical support, Sara Pérez-Jaume for her assistance in formal analysis and data curation and the Genomics Unit at the CRG for assistance with the sequencing. The project also had the support from the Asociación Pablo Ugarte (APU). We are grateful to members of the Mora and Cayetano González laboratories for helpful discussions, and to the Band of Parents at Hospital Sant Joan de Déu for supporting the overall research activities of the Developmental Tumor Laboratory, PCCB. This study is dedicated to the memory of all Ewing sarcoma patients at HSJD who have succumbed to the disease and whose families have devoted their efforts to understand the origin of the tumor.

## ETHICS DECLARATION

Animal experiments were conducted following protocols approved by the Institutional Ethics Committee on Experimental Animals, in full compliance with Spanish and European laws and regulations. Approval to conduct this study was obtained from the ethics committees on Clinical Research in accordance with the guidelines of the Helsinki Declaration (1975), revised in 1983.

The authors declare no potential conflicts of interest.

## ADDITIONAL INFORMATION

This work was funded by La Marató TV3 Foundation (201928) and received the support from the grant PI22/00364 (funded by Instituto de Salud Carlos III (ISCIII) and co-funded by the European Union) and the “Xarxa de Bancs de tumors” sponsored by Pla Director d’Oncologia de Catalunya (XBTC).

---

**Figure S1.**
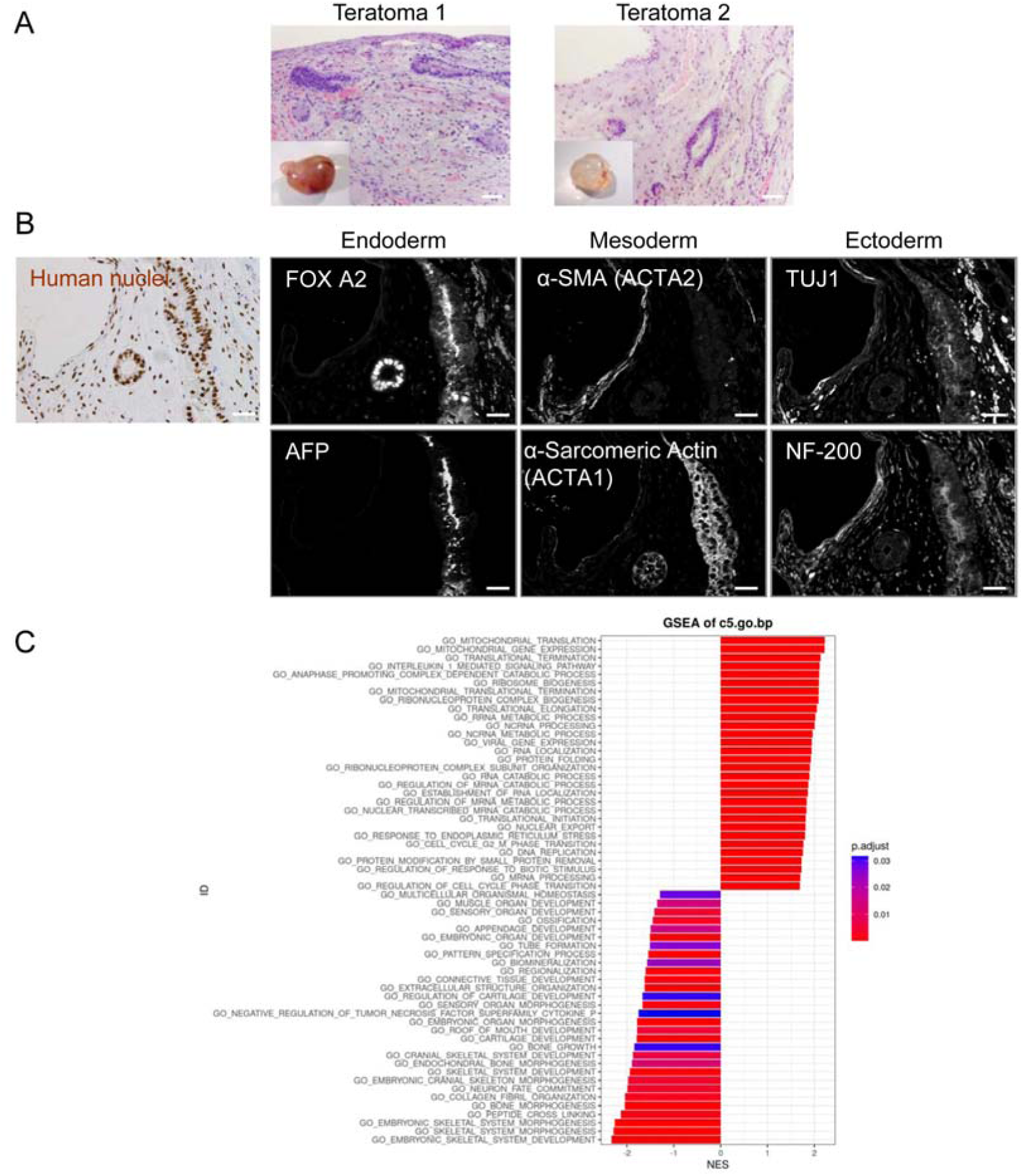
Formation of Teratomas from heSCs. (A) Gross morphology and hematoxylin and eosin (H&E) staining of two teratomas from heSCs xenografted into testis of immunosuppressed mice. Scale bars, 200μm. (B) Immunofluorescent labeling of structures corresponding to the three main embryonic germ layers in teratomas. On the left: human nuclei detected by immunohistochemistry (IHC) in serial sections. Scale bars, 100μm. (C) Barplot of significantly enriched terms (adj. P val. <0.05) in control heMSC-1 cells (normalized enrichment score, or NES, positive) vs. hpMSCs (NES negative).

**Figure S2.**
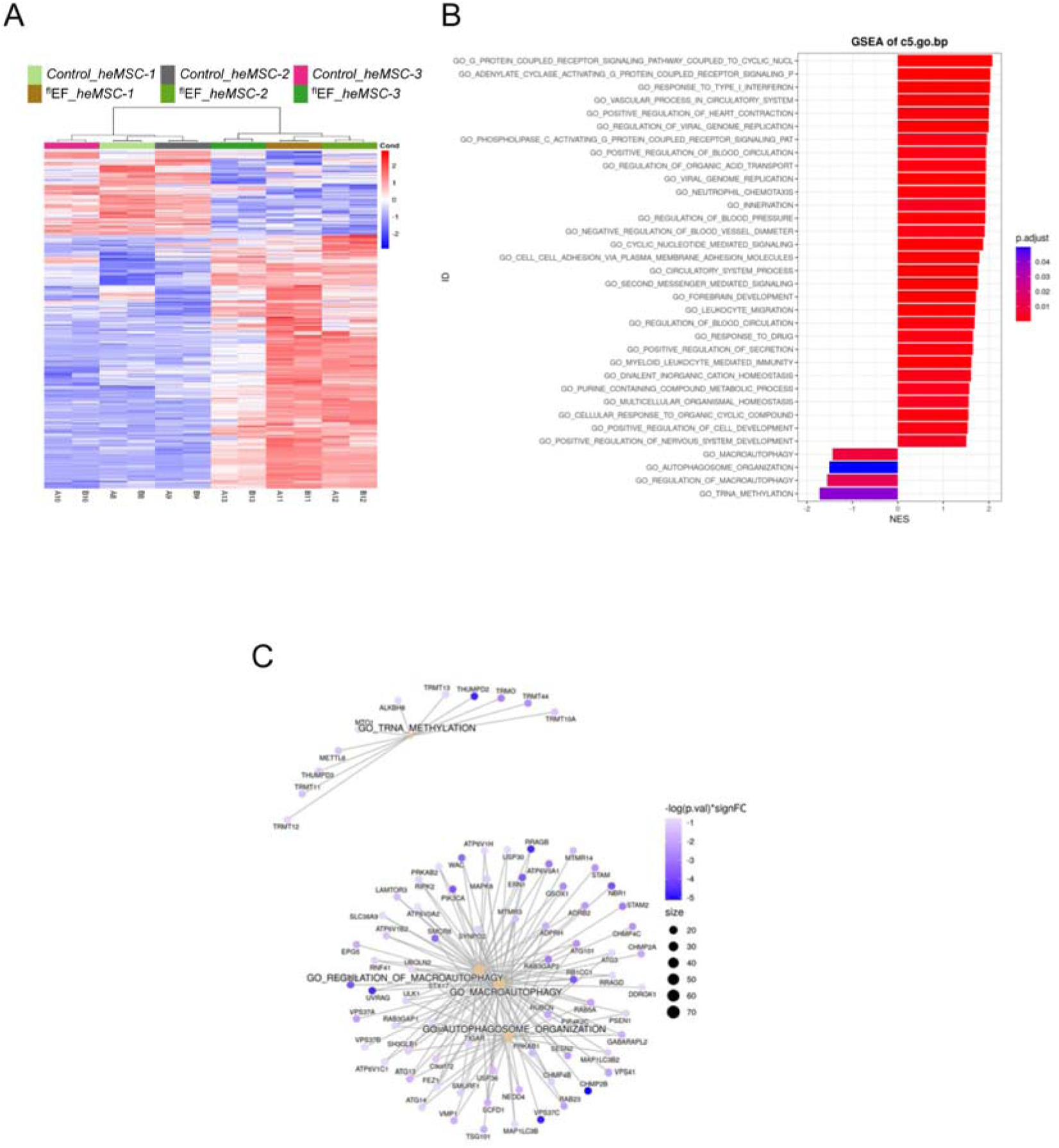
EWS-FLI1 Expression in Teratoma-derived heMSCs. (A) Heatmap of differential expressed genes (adj. P val. <0,05; |logFC|>1) in heMSCs infected with EWS-FLI1 when compared to heMSCs infected with empty vector. (B) Barplot of significantly enriched terms in EF-heMSCs (NES, positive) vs. control heMSCs (NES negative). (C) Gene-concept networks of top 5 significantly underrepresented terms in EF-heMSCs vs. control heMSCs.

**Figure S3.**
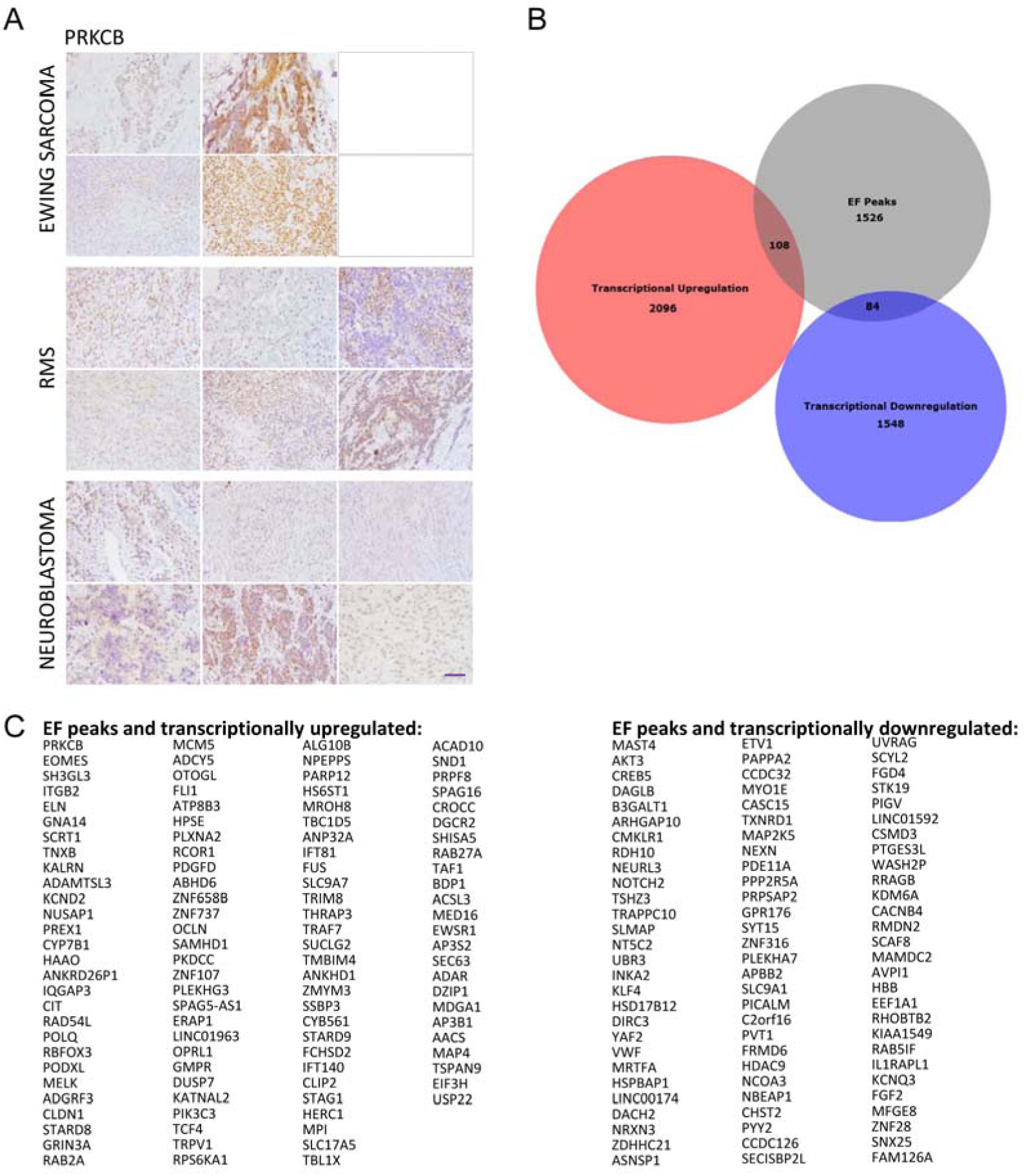
Identification of Direct EWS-FLI1 Targets with Transcriptional Changes in heMSCs. (A) PRKCB expression in Ewing sarcoma, rhabdomyosarcoma and neuroblastoma tumor samples, detected by IHC. Magnification bar, 50µm. PRKCB expression in two of the Ewing sarcoma tumors could not be assessed due to the deleterious effect of decalcification on the PRKCB antigen. (B, C) Venn diagram and genes associated to EWS-FLI1 bound regions (grey) were overlapped with overexpressed (red) or downregulated (blue) genes in heMSCs.

**Figure S4.**
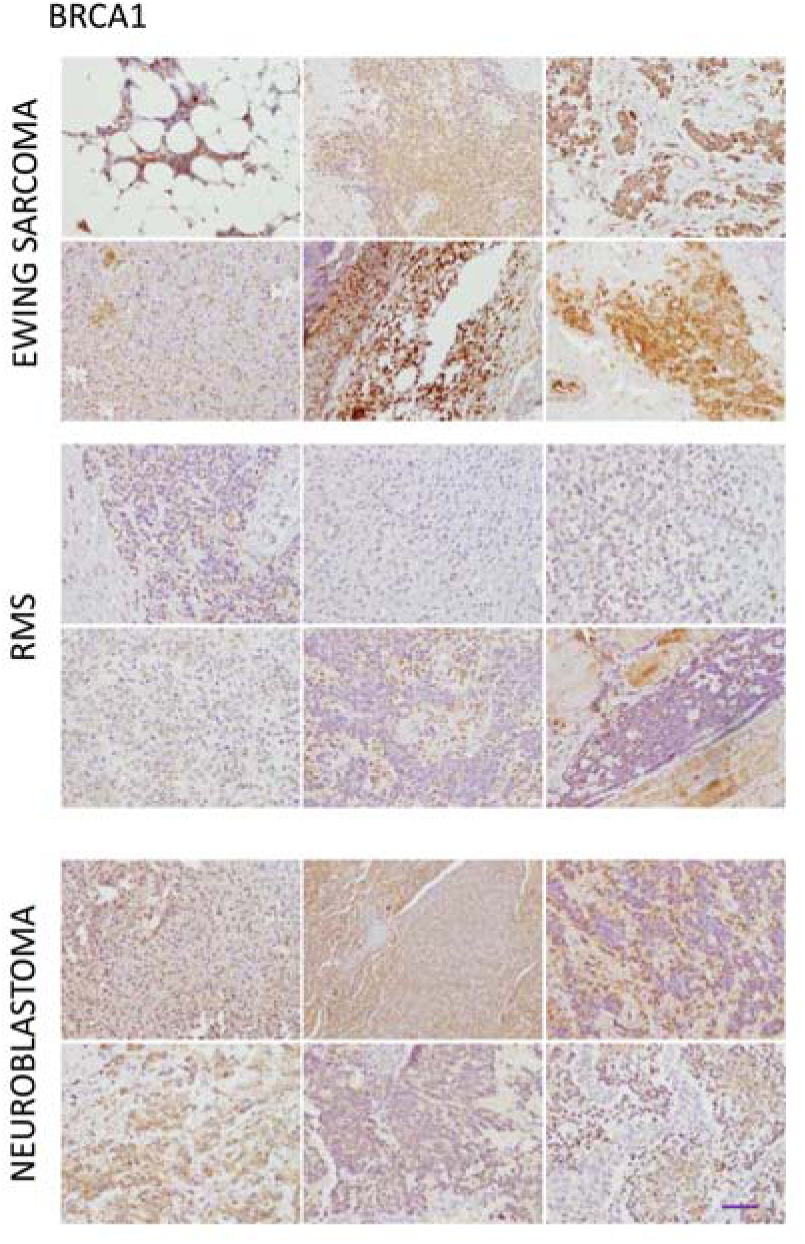
BRCA1 Expression in Developmental Tumors. BRCA1 expression in Ewing sarcoma, rhabdomyosarcoma and neuroblastoma tumor samples, detected by IHC. Magnification bar, 50µm.

**Figure S5.**
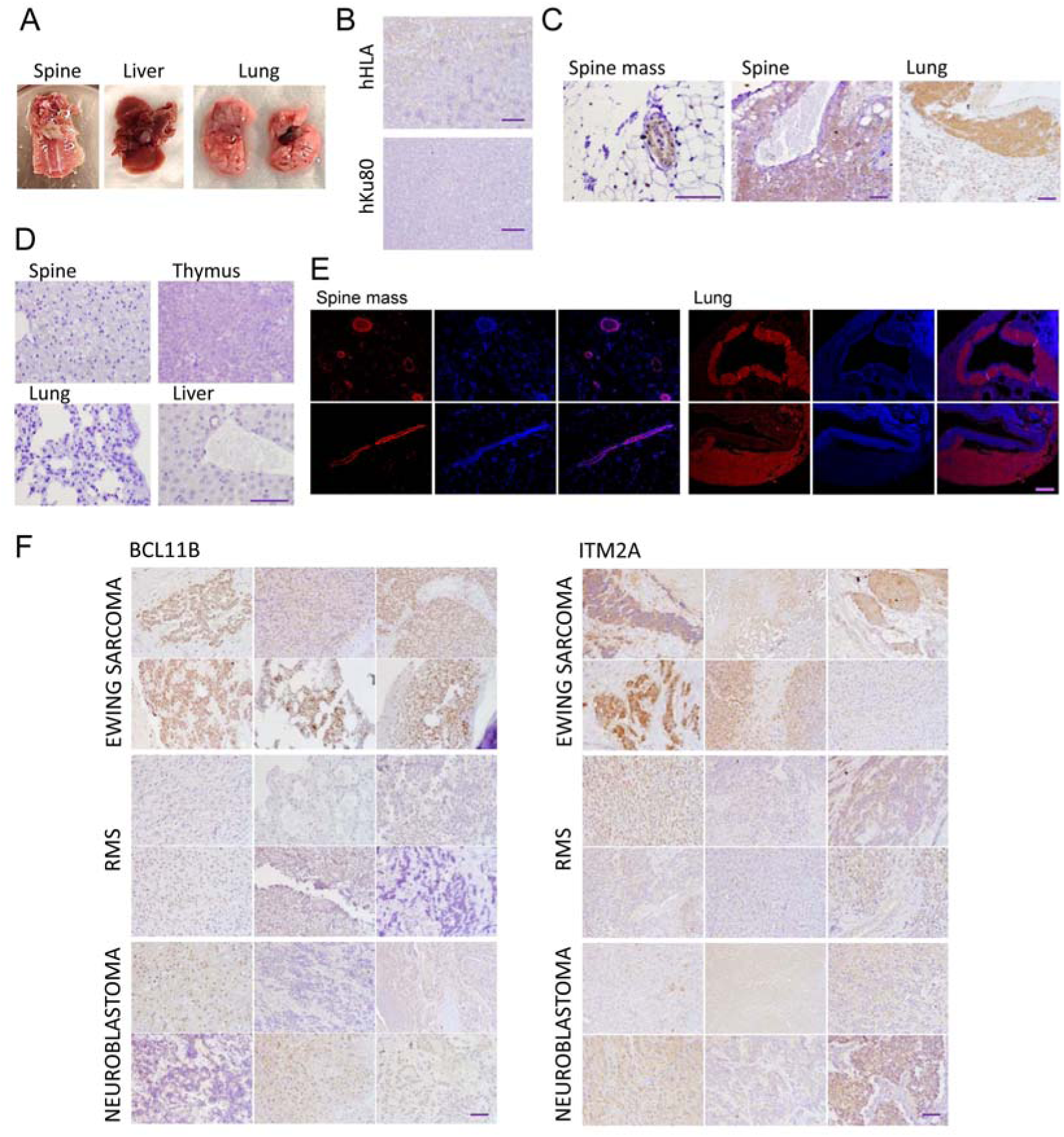
EF-heMSCs Form Tumors in Vivo. (A) Gross morphology of lesions generated upon injection of EF-infected heMSC lines. (B) IHC of mouse thymus to assess the specificity of hHLA and hKu80 antibodies. Scale bars, 50µm. (C) IHC to detect Flag expression in lesions from NOD/SCID mice injected with EF-infected heMSC lines. Scale bars, 50µm. (D) IHC of different mouse tissues to assess the specificity of the Flag antibody. Scale bars, 50µm.G (E) Immunofluorescent detection of Flag expression (red) in cells of EF-heMSC-derived lesions, with cell nuclei distribution identified by DAPI staining (blue). Two different sections are shown. (F) Expression of the ESFT markers BCL11B and ITM2A in developmental tumors, assessed by IHC. Scale bars, 50µm.

